# SynGAP forms biocondensates at sub-micromolar concentrations and recruits PSD95 and receptor oligomers, functioning as a key initiator of PSD formation

**DOI:** 10.1101/2025.04.22.649955

**Authors:** Saahil Acharya, Taka A. Tsunoyama, Christian Hoffmann, Perez Gerard Aguilar, Irina Meshcheryakova, Aya Nakamura-Norimoto, Tara Mastro, Ward G. Walkup, Takahiro K. Fujiwara, Mary B. Kennedy, Dragomir Milovanovic, Akihiro Kusumi

**Affiliations:** Membrane Cooperativity Unit, Okinawa Institute of Science and Technology Graduate University, Onna-son, Okinawa 904-0495, Japan; Laboratory of Molecular Neuroscience, German Center for Neurodegenerative Diseases (DZNE), 10117 Berlin, Germany; Division of Biology and Biological Engineering, California Institute of Technology, Pasadena 91125, California, U.S.A.; Institute for Integrated Cell-Material Sciences (WPI-iCeMS), Kyoto University, Kyoto 606-8501, Japan

## Abstract

A key issue in neuronal circuit regulation is how synapse formation is initiated. Synapse formation could start when one or more synaptic scaffold proteins that can initiate synapse formation reach certain threshold concentrations in the dendritic shaft, which might lead to their oligomerization or even liquid-liquid phase separation (LLPS). By combining in vitro reconstitution of purified proteins with live-cell single-molecule and confocal imaging, we demonstrated that SynGAP alone forms assemblies of nanoscale clusters containing several to several tens of molecules at 10-nM order concentrations and micron-scale LLPS hydrogel-like condensates at submicromolar concentrations. The trimers of SynGAP’s intrinsically disordered region (IDR) induced by its coiled-coil domain are responsible for SynGAP condensation. CaMKII-mediated phosphorylation moderately suppresses SynGAP condensation, and also increases condensate liquidity. While PSD95 fails to form assemblies under these conditions, it is recruited to SynGAP condensates by specifically binding to the PDZ-binding motif of SynGAP. SynGAP[PSD95] condensates selectively immobilize postsynaptic transmembrane proteins, Neuroligin1 and AMPAR-TARP2 complexes, in a manner dependent on their oligomerization state, indicating cooperative recruitment dynamics among SynGAP, PSD95, and transmembrane components, which might mimic initial PSD assembly. These findings suggest that SynGAP may act as a primary nucleator of postsynaptic density assembly, challenging the PSD95-centered models.

## INTRODUCTION

Mechanisms underlying formation of the post-synapse have been extensively investigated. Südhof (2021) proposed that synapse formation is initiated by the clustering of synaptic adhesion molecules, specifically clustering of postsynaptic Neuroligin1 (Nlg1) associating with presynaptic Neurexin1β (Nxn). He proposed that these clusters become the “nucleus” for assembling postsynaptic scaffold proteins, notably including PSD95 which then recruits α-amino-3-hydroxyl-5-methyl-4-isoxazole-propionate-type glutamate receptors (AMPARs) by binding to TARPs (transmembrane AMPA receptor-associated proteins), N-methyl-D-aspartate receptors (NMDARs), and additional scaffold proteins, such as Shank and Homer, which link the postsynaptic density (PSD, Kennedy 2000) to the actin cytoskeleton (Mondin et al., 2011, Chen et al., 2023; Dosemeci et al., 2016; Shiraishi-Yamaguchi et al., 2007; Valtschanoff et al., 2001). Thus, PSD95 clustering is a key step in the early formation of the PSD. However, the precise mechanisms by which initial small PSD95 assemblies associate with neuroligin clusters, TARPs, AMPARs, and NMDARs; and eventually grow into the sub-micron-scale assemblies that constitute the mature postsynaptic density (PSD) are still not fully understood.

Localization of PSD95 near the plasma membrane (PM) is mediated through N-terminal palmitoylation (Sturgill et al., 2009), with subsequent multimerization (Christopherson et al., 2003). Recent reports suggest that liquid-liquid phase separation (LLPS), also referred to as “condensate formation” plays an important role in further clustering of PSD95 and formation of the PSD. These reports include *in-vitro* observations that PSD95 combines with partial sequences of other PSD proteins, including SynGAP, GKAP, Shank3, and/or the cytoplasmic tail domains of GluN2B and TARP2, to form condensates (Zeng et al., 2016; Zeng et al., 2018; Zeng et al., 2019; Shen et al., 2023; Zhu et al., 2024). In some *in-vitro* studies, PSD-protein condensates centered around PSD95 formed on supported membrane lipid bilayers induced clustering of the cytoplasmic domains of GluN2B, a subunit of the NMDAR, and selectively enriched a small fragment of SynGAP comprised of the coiled-coil domain and PDZ-binding motif (PBM) (Zeng et al., 2018).

SynGAP was initially identified as a GTPase activating protein (GAP) isolated from the forebrain PSD fraction and located in the PSD *in vivo* (Chen et al., 1998; Kim et al., 1998; Krapivinsky et al., 2004). It has been implicated in modulating spine formation and synaptic strength through its GAP activity and/or its interaction with PSD95 (Vazquez et al., 2004; McMahon et al., 2012, Komiyama et al., 2002; Rumbaugh et al., 2006; Wang et al., 2013). SynGAP undergoes phosphorylation-induced displacement from the PSD, where it can compete with TARP/AMPARs for binding to PSD95, acting as a negative regulator of excitatory synaptic strength (Vazquez et al., 2004; McMahon et al., 2012; Araki et al., 2015; Walkup et al., 2016). Further investigations revealed that SynGAP in fact plays a significant structural role that is independent of its enzyme activity (Walkup et al., 2016; Araki et al., 2024).

SynGAP is present at roughly equivalent copy numbers to PSD95 in the PSD (≈300 copies; Cheng et al., 2006; Sugiyama et al., 2005; Chen et al., 2005), yielding an equivalent concentration of ≈5-25 µM in spines (assuming the presence of a single PSD per spine and a spine volume of 0.02-0.1 µm^3^; Leuner et al., 2010; Arellano et al., 2007). Despite this abundance, the significance of SynGAP in PSD assembly has been considered secondary to that of PSD95 and CaMKII, perhaps due to its negative regulatory role in spine formation (Vazquez et al., 2004; Araki et al., 2015). Like PSD95, CaMKII is regarded as a principal nucleator within the PSD. Indeed, activated CaMKII combines with the dimerized carboxyl tails of GluN2B (Bayer et al., 2006) and undergoes condensation by LLPS *in-vitro* (Hosokawa et al., 2021). In the presence of PSD-95, it forms a phase-in-phase condensate in which PSD-95 occupies the core region (Hosokawa et al. 2021). However, in-vitro studies examining condensates with PSD95 and CaMKII have exclusively used mixtures with various other PSD-protein fragments (Zeng et al., 2016; Zeng et al., 2018; Zeng et al., 2019; Hosokawa et al. 2021; Shen et al., 2023; Zhu et al., 2024), which has left the individual contributions of each protein and protein fragment to condensation ambiguous. Moreover, the protein concentrations used in these studies were generally above 1 µM and frequently exceeded 10 µM, which might be comparable to the concentrations in mature spines, yet is considerably higher than those in emerging postsynaptic sites.

None of these prior *in-vitro* studies, addressed the formation of condensates by SynGAP because they employed only a truncated fragment of SynGAP comprising the coiled-coil domain and PDZ-binding motif (PBM) — a mere 162 amino-acid sequence out of the full length 1,308 amino-acid full-length SynGAPαI (Zeng et al., 2016; Zeng et al., 2018; Zeng et al., 2019; Zhu et al., 2024). We note that the disorder scores predicted by ANCHOR2 (Meszaros et al., 2018) suggest that SynGAP contains a large, 409 amino-acid intrinsically disordered region (IDR) between the C-terminus of its RasGAP domain and the coiled/coil domain. The IDR makes up ≈30% of the full-length protein (Gamache et al., 2020). Because IDRs have often been implicated in formation of condensates by LLPS (Turoverov et al., 2019), we investigated whether full-length SynGAP alone forms LLPS-induced condensates, contributing to the assembly of the other key PSD proteins with which it associates, including PSD95. In particular, we have examined whether SynGAP condensation occurs at sub-micromolar concentrations, which would be more characteristic of a nascent postsynaptic region as the PSD begins to form.

Here we use full-length proteins to demonstrate that SynGAP alone, unlike PSD95 alone, forms hydrogel-like condensates by LLPS at low micromolar concentrations *in-vitro* and at sub-micromolar concentrations in living cells. We attribute the propensity of SynGAP to undergo LLPS to its large IDR in tandem with the coiled-coil domains which form trimers (Zeng et al., 2016). Thus, SynGAP trimerization of the IDRs via the coiled-coil domains drives SynGAP to undergo LLPS.

PSD95 is recruited to these SynGAP condensates via binding of its PDZ domains to the PDZ ligand of SynGAP (PBM). The SynGAP/PD-95 condensates are particularly effective in recruiting oligomers of full-length transmembrane postsynaptic proteins, such as Nlg1 and TARPs (oligomerized via binding to AMPARs). We propose that SynGAP condensation may serve as a seminal event in initiating assembly of the PSD by orchestrating the recruitment of PSD-95 and transmembrane postsynaptic receptors, thereby establishing a fundamental scaffold for the subsequent growth and maturation of PSD assemblies.

## RESULTS

### SynGAP forms liquid-liquid phase-separated condensates at low micromolar concentrations, and recruits PSD95 as a client, *in-vitro*

We first purified two proteins, SynGAPα1’s partial sequence containing the IDR-CC-linker-PBM tagged with mGFP (mGFP-SynGAPα1-IDR-CC[PBM]) and PSD95 tagged with mScarlet (PSD95-mScarlet) (**Figure 1B; Supplementary Figure 1**), and examined their abilities to undergo LLPS to form condensates *in-vitro* (whenever the meanings are clear, we will simply call these proteins SynGAP and PSD95, respectively). We varied the concentrations of SynGAP and PSD95 in the range of 1∼7.6 µM and that of the crowding reagent PEG8000 in the range of 0∼3%, and generated phase diagrams for SynGAP and PSD95 (**Figure 1C**). The results demonstrated that SynGAP forms condensates at concentrations of ≥2 µM with ≥1% PEG8000, whereas the full-length PSD95 alone forms condensates at concentrations of ≥4 µM with ≥3% PEG8000. Thus, SynGAP has a propensity to form condensates at lower concentrations compared to PSD95. Consistently, 0.5 and 2 µM concentrations of PSD95 (at 1% PEG8000), which cannot form condensates, can form co-condensates with SynGAP in the presence of 2 µM SynGAP (**Figure 1D**).

**Figure 1.**
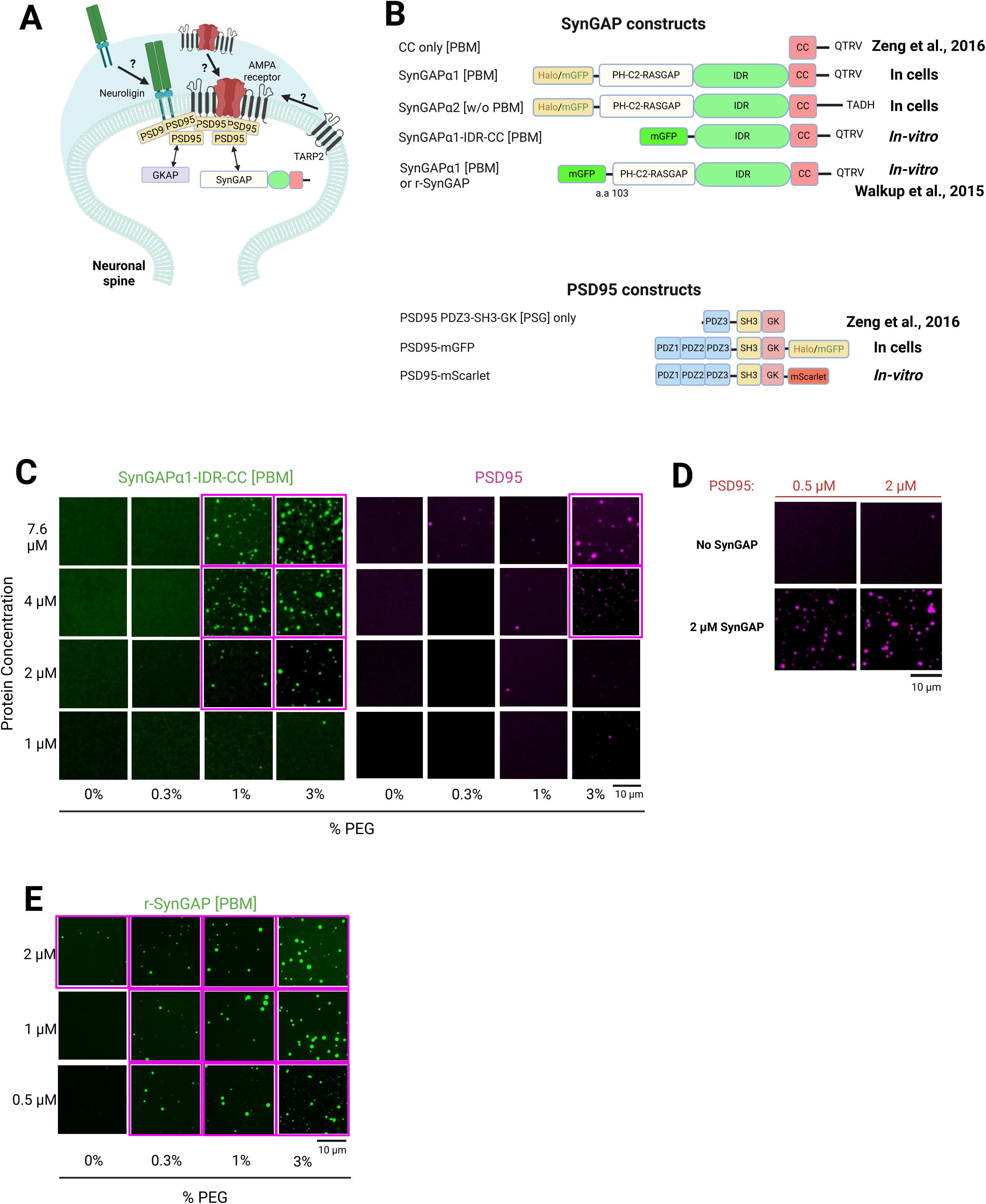
*In-vitro*, SynGAP alone can form liquid-liquid phase-separated condensates. (A) Schematic of the neuronal post-synaptic spine, showing some of the key molecules interacting in the synapse. PSD95, an essential PSD scaffold protein, was proposed to be the LLPS driver for PSD assembly. GKAP and SynGAP are cytoplasmic proteins interacting with PSD95 and TARPs, and Neuroligins are transmembrane cell-cell adhesion proteins that can bind to PSD95. AMPA receptors bind to PSD95 by way of TARPs. The biophysical mechanisms by which PSD95 and its binding molecules are assembled and retained in the post synapse, even when their bulk concentrations are low (e.g., during the post-synapse formation), remain to be discovered. (B) The SynGAP and PSD95 constructs employed in the present research. The SynGAP construct used by Zeng et al. (2016) and subsequent works is also shown. Full-length isoforms of SynGAPα1 and α2 differ in that the α1 isoform contains the C-terminal PDZ binding motif (PBM), but the α2 isoform lacks the PBM. The r-synGAP construct, containing almost the entire PH-C2-RasGAP domain in addition to the IDR, CC, and C-terminal domain, missing only the first 103 amino acids (α1 isoform; Walkup et al., 2015), was used for some *in-vitro* experiments. (C) Coarse phase diagrams for the *in-vitro* condensation of mGFP-SynGAPα1-IDR-CC[PBM] (left) and PSD95-mScarlett (right) with various concentrations of proteins and PEG8000 (w/v) at physiological salt concentrations (150 mM NaCl), observed 2 min after PEG addition. (D) PSD95-mScarlet below 2 µM does not form condensates, whereas it colocalizes with the mGFP-SynGAP condensates when SynGAP coexists at 2 µM (with 1% w/v PEG 8000, 150 mM NaCl). (E) Coarse phase diagram for the *in-vitro* condensation of (mGFP tagged) r-SynGAP with various concentrations of proteins and PEG8000 (w/v) at physiological salt concentrations (150 mM NaCl), observed 2 min after PEG addition.

When we examined purified r-SynGAP, which lacks only the initial 103 N-terminal amino acids (Walkup et al., 2015) with mGFP tag (**Figure 1A**) in the same way, its condensation ability was even higher. r-SynGAP forms condensates at concentrations of ≥0.5 µM with ≥0.3% PEG (**Figure 1E**). This could be due to interactions between N-terminal domains of r-SynGAP, such as inter-molecular C2-RasGAP domain interactions (intra-molecular C2-RasGAP interactions have been shown; Pena et al., 2008).

We also used r-SynGAP to quantitatively assess its effects on and interactions with PSD95. r-SynGAP, PSD95, or their mixtures in solution were allowed to form condensates, and after separating the condensates by centrifugation, the amounts of r-SynGAP and PSD95 remaining in solution or in the condensates were evaluated by SDS polyacrylamide gel electrophoresis. In the experiments employing r-synGAP alone and PSD95 alone, 23% of SynGAP and only 1% of PSD95 were found in the pellet fraction, consistent with the phase diagrams shown in **Figure 1C**. In the experiments mixing equimolar concentrations of SynGAP and PSD95, 59% and 33% of these proteins, respectively, were found in the pellet fraction (**Supplementary Figure 1F**). These results are partially consistent with a previous biochemical observation using cells expressing these proteins (Araki et al., 2020), showing that PSD95 can enhance the formation of SynGAP condensates.

### SynGAP forms condensates in cells at sub-micromolar concentrations and recruits PSD95 as a client

We examined the condensate formation of full-length SynGAP (mGFP-conjugated) and PSD95 (mGFP-conjugated) separately expressed in L cells, using confocal microscopy in the confocal plane at 1 µm above the coverslip. To quantitatively evaluate condensate formation, we always measured the cytoplasmic concentration of expressed mGFP-linked proteins. The signal intensity in each pixel was calibrated by the signal intensities of known concentrations of mGFP in solution (**Supplementary Figure 2A**), and the amount of the examined protein in condensates in the confocal plane was evaluated. The extent of condensation (clustering) was evaluated using the clustering coefficient, which is the total fluorescence intensity in the detectable condensates normalized by the cell area measured in the confocal image, and thus is proportional to the expected protein concentration in the cytoplasm if the protein in the condensates were dispersed throughout the cytoplasm (see Materials and methods). The clustering coefficient was plotted as a function of the total cytoplasmic concentration averaged over the cell (**Figures 2A and 2B**).

**Figure 2.**
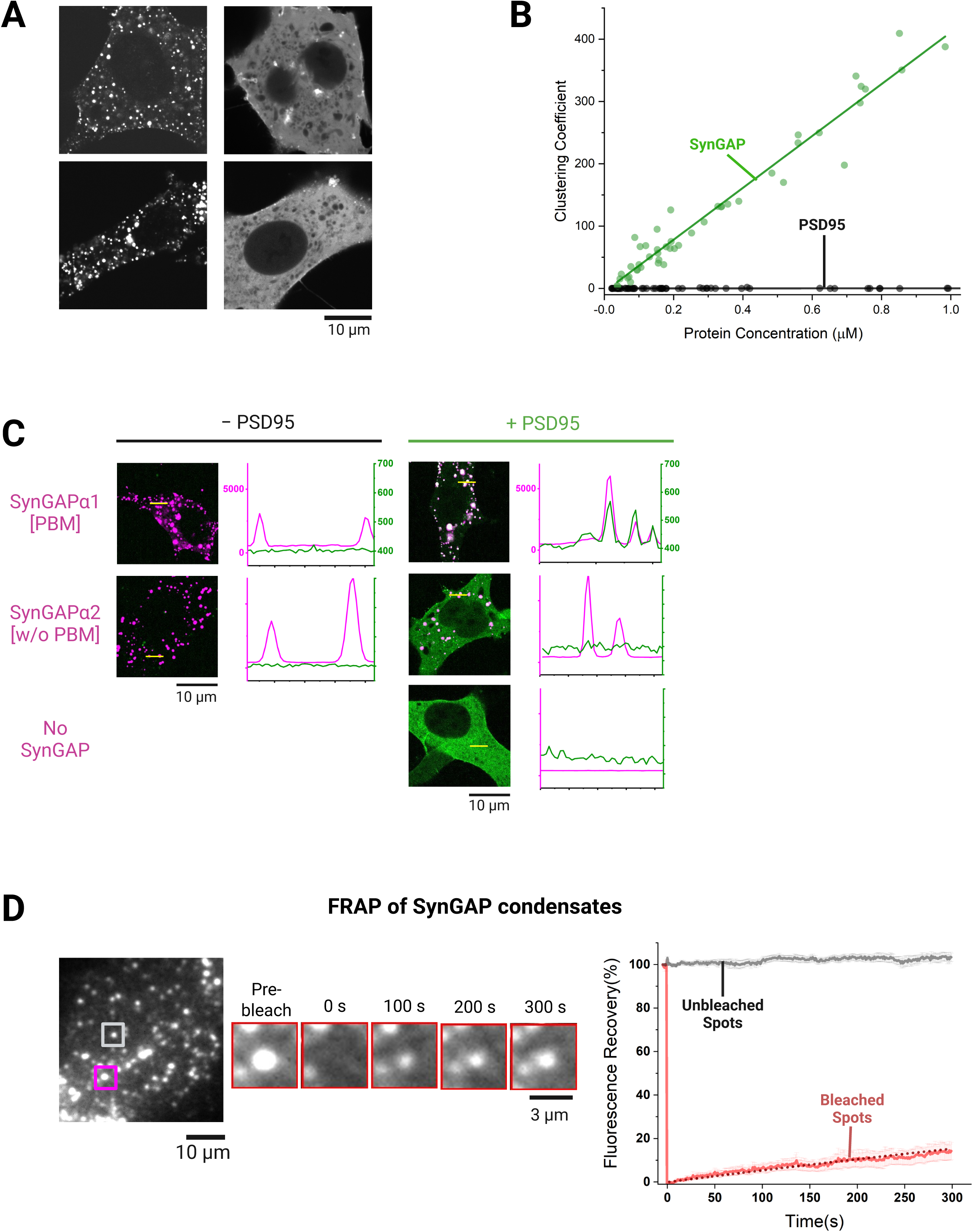
In cells, SynGAP alone (at >0.04 µM), but not PSD95 alone (even at ≈1 µM), forms LLPS-induced condensates and recruits PSD95 as a client. (A) Typical fluorescence images of L cells expressing mGFP-SynGAPα1 (full length) alone (left) and PSD95-mScarlet alone (right) at 1 µM, showing that SynGAP, but not PSD95, forms intracellular condensates under these conditions. (B) In the L-cell cytoplasm, mGFP-SynGAPα1 (full length) alone, but not PSD95-mGFP alone, forms condensates at sub-micromolar concentrations. mGFP-SynGAP condensates are found at >0.04 µM. The clustering coefficient is plotted as a function of the proteins’ average cytoplasmic concentrations in each cell. Each data point represents the result from a single cell (47 cells for mGFP-SynGAPα1 and 54 cells for PSD95-mGFP; three independent replicates). Lines indicate the results of linear fitting for visualization. (C) PSD95 is recruited to SynGAP condensates as a client by binding to the PBM of SynGAP. (Left) In L cells, both Halo-SynGAP isoforms α1 (containing PBM) and α2 (lacking PBM) form condensates in the absence of PSD95. (Right) PSD95-mGFP did not exhibit colocalization with the condensates of the α2 isoform, which lacks PBM, but colocalized with the condensates of the α1 isoform, which contains PBM, unequivocally demonstrating that PSD95 is recruited to the α1 condensates as a client by binding to the PBM of the α1 isoform. Line graphs (magenta, SynGAP; PSD, green) indicate the pixel intensities along the yellow lines shown in the images on the left. (D) FRAP of mGFP-SynGAPα1 (full length) condensates in live L cells expressing ≈1 µM mGFP-SynGAPα1 in the cytoplasm, showing the hydrogel-like nature of the SynGAP condensate. (Left) A representative cell image and a typical FRAP time course of a condensate, indicated by the magenta square in the cell image. The condensate indicated by the grey square in the cell image was used as a reference to evaluate the photobleaching effect during the FRAP observation. (Right) FRAP recovery results for the bleached (red) and unbleached (black) spots (mean ± SE from 6 and 17 condensates) in 6 and 17 cells for bleached and unbleached condensates, respectively, from three independent replicates. Dotted line indicates the fitting result using a single exponential recovery function.

The results showed the marked difference between full-length SynGAP and PSD95. With an increase of the total protein concentration in the cytoplasm (x axis of **Figure 2B**) upon formation of protein condensates, larger amounts of the expressed protein would be incorporated in the condensates without increasing the cytoplasmic protein concentration outside the condensates, and thus the clustering coefficient should increase linearly with the total averaged concentration. SynGAP exactly showed this tendency, and the graph revealed that SynGAP condensation starts occurring at ≈0.04 µM. In contrast, no indication of condensation was found for PSD95 below 1 µM.

Next, we examined the possibility of co-condensation of SynGAP and PSD95 at submicromolar concentrations in live cells. For this purpose, we employed two SynGAP isoforms (tagged with Halo at their N termini), which can and cannot bind to PSD95 (**Figure 1B**): SynGAPα1, containing the C-terminal PDZ binding motif (PBM) of QTRV, which allows PSD95 binding, and SynGAPα2, an isoform that differs from SynGAPα1 only in the C-terminal region, where it lacks the PBM and thus cannot bind PSD95. Both SynGAP isoforms formed condensates at an average cytoplasmic concentration of ≈1 µM (**Figure 2C, left column**). When PSD95 (mGFP-conjugated) was co-expressed, PSD95 colocalized with the condensates of the SynGAPα1 isoform, which contains the PBM (with a concomitant large reduction of the cytoplasmic PSD95 concentrations), but not with the condensates of the SynGAPα2 isoform, which lacks the PSD95-binding motif (**Figure 2C, right column**; also see **Supplementary Figures 2B and 2C**). This result unequivocally demonstrates that PSD95 does not undergo co-condensation with SynGAP, but is simply recruited to the SynGAPα1 condensates as a client by specifically binding to its PBM (the L cells employed here do not express PSD95 at levels detectable by western blot; **Supplementary Figure 2D**).

Fluorescence recovery after photobleaching (FRAP) experiments showed a recovery half-life (τ_1/2_) of 21 min, which was obtained by fitting the recovery data with a single exponential function assuming full recovery (direct FRAP data showed that ≈14% of the SynGAP molecules in the condensates exchange with those located in the bulk cytoplasm in 5 min; **Figure 2D**), indicating that SynGAP condensates are hydrogel-like. In conclusion, SynGAP forms hydrogel-like condensates by undergoing LLPS at submicromolar concentrations in the live-cell cytoplasmic environment, and recruits PSD95 as a client via its C-terminal PBM (**Figure 2**), while PSD95 binding to SynGAP enhances SynGAP condensation (**Figures 1E and 1F**).

### SynGAP’s IDR linked to CC domain is responsible for SynGAP condensation

We further investigated which domains of SynGAP are responsible for condensation by performing quantitative examinations. For this examination, we varied the average protein concentration in the cytoplasm in the range between 0 and 1.1 µM, and examined the formation of condensates, using a clustering coefficient. First, we found both full-length SynGAP isoforms form condensates in the concentration range between 0.07 and 1.1 µM. The clustering coefficient linearly increases with an increase of their average cytoplasmic concentrations from 0.07 to 1.1 µM (**Figure 3A**). This indicates that when the average cytoplasmic concentration of SynGAP becomes higher than 0.07 µM, SynGAP molecules exceeding this concentration are incorporated in SynGAP condensates.

**Figure 3.**
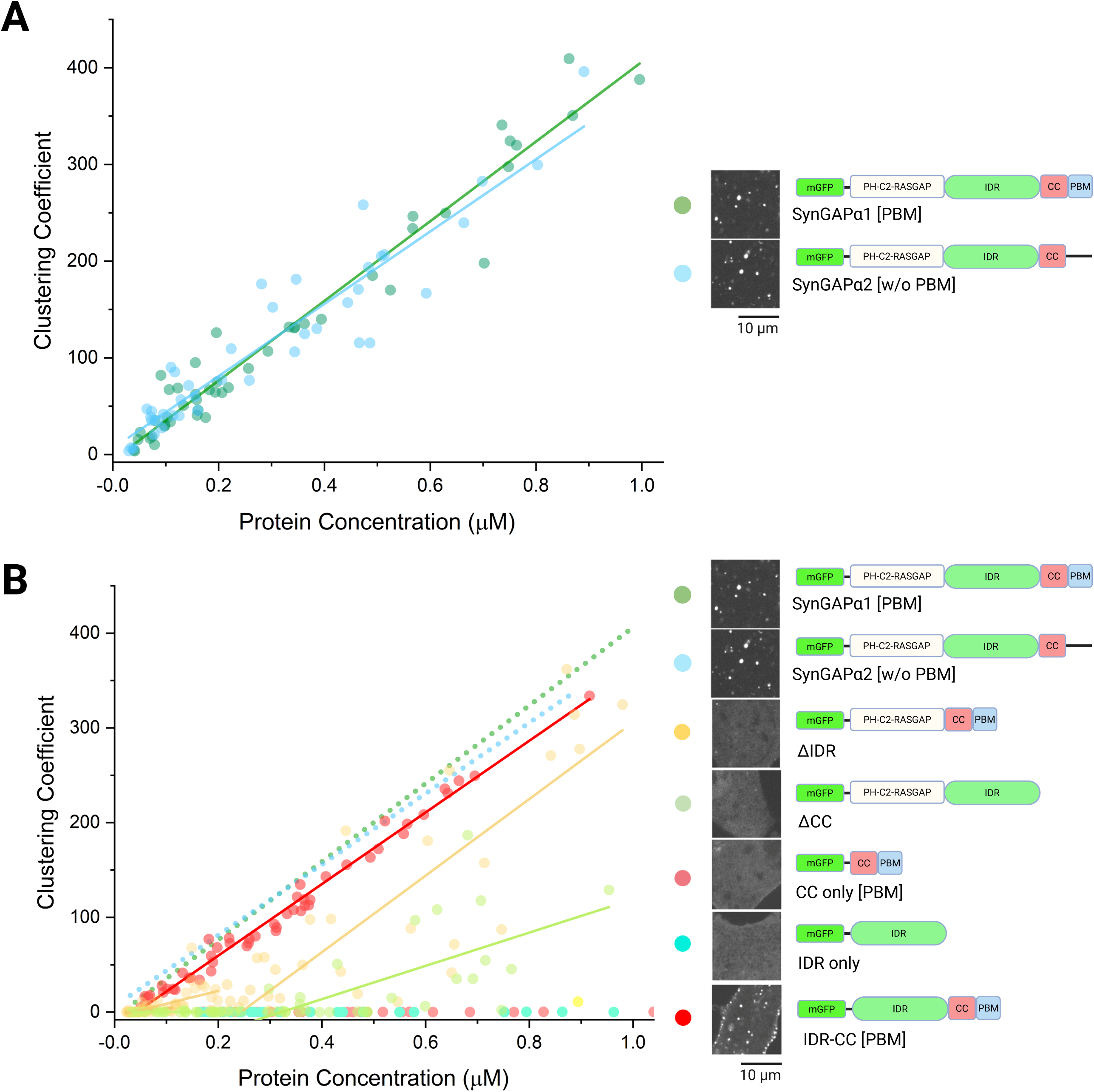
SynGAP’s IDR linked to the CC domain is responsible for generating SynGAP’s LLPS condensates, while CaMKII-induced SynGAP phosphorylation moderately suppresses condensation. (A) Condensate formation of mGFP-tagged SynGAP isoforms (right column) expressed in L cells, examined as a function of the total cytoplasmic protein concentration in each cell. Linear fitting was performed to facilitate visualization in the entire range. The numbers of examined cells: 47 for SynGAPα1 and 40 for SynGAPα2. The middle column shows typical images of cells expressing the examined constructs. (B) Condensate formation of mGFP-tagged SynGAP constructs (right column) expressed in L cells, examined as a function of the total cytoplasmic protein concentration in each cell. Linear fitting was performed to facilitate visualization, in the range of 0.3-1.1 µM for all constructs. The linear fits shown in **A** for SynGAPα1 and SynGAPα2 are shown again (dotted lines, without data points) for reference. The numbers of examined cells: 89 for ΔIDR, 49 for ΔCC[PBM], 58 for CC [PBM]only, 43 for IDR only, and 50 for IDR-CC [PBM]. The middle column shows typical images of cells expressing the examined constructs.

Next, we examined various partial sequences of SynGAP for condensate formation as a function of the average cytoplasmic concentration (**Figure 3B**). The mutant lacking IDR (ΔIDR) exhibited a lower ability to form condensates, and the mutant lacking the CC domain (ΔCC), which induces SynGAP trimerization (Zeng et al., 2016), showed even lower ability (Their slopes at higher concentrations are somewhat different from those for the full-length SynGAP isoforms. The reason for this difference is unknown, but it might be due to the differences in the condensate sizes, which would affect the detectability by confocal microscopy). The CC domain alone, which was previously used as the “SynGAP construct” (Zeng et al., 2016), and the IDR alone did not form condensates in this concentration range.

Remarkably, when the IDR was linked to the CC domain, the linked construct behaved almost identically to the full-length SynGAP isoforms. These results demonstrated that the CC-domain-induced IDR trimers or SynGAP trimers are the basic unit for the formation of SynGAP condensates.

### SynGAP forms nano-scale clusters in cells at nanomolar concentrations

While we found micron-scale condensates of SynGAP start appearing at sub-micromolar concentrations, we suspected that at the lower concentration tail regime, smaller nano-scale clusters, which would act as nucleation sites for condensation, might appear. To detect nano-scale condensates, we observed the bottom PM using TIRF laser illumination so that we can detect immobilized or slowly diffusing nano-clusters there because nano-clusters diffusing rapidly in the cytoplasm would be difficult to detect. The laser illumination intensity was adjusted so that SynGAP clusters consisting of four to ≈250 molecules become detectable. Full-length SynGAP (Halo-SynGAP) was observed at the average cytoplasmic concentrations ranging from 10 to 300 nM. The cytoplasmic SynGAP concentrations were determined by observing the same cells by confocal microscopy.

SynGAP forms diffraction-limited nano-scale clusters at cytoplasmic concentrations of 10 nM and above, while PSD95 hardly forms any such clusters in the concentrations up to 270 nM (**Figure 4A**). SynGAP shows a concentration dependent increase in the density of clusters on the basal PM up to ∼ 100 nM, which then plateaus with increasing concentration at a number density of 0.14 ± 0.02 clusters/µm^2^ (obtained by fitting with asymptotic function) (**Figure 4B**). The plateau might occur because, as SynGAP clusters grow, they tend to detach from the PM to move into deeper cytoplasm. Meanwhile, in the overall cell system, the copy number of SynGAP molecules incorporated into clusters would linearly increase with concentration, forming micron-scale clusters (**Figure 2B and Figure 3A**). Meanwhile PSD95 does not form detectable nano-scale clusters on the basal PM in the range between 10 and 300 nM (**Figure 4A and Figure 4B**).

**Figure 4.**
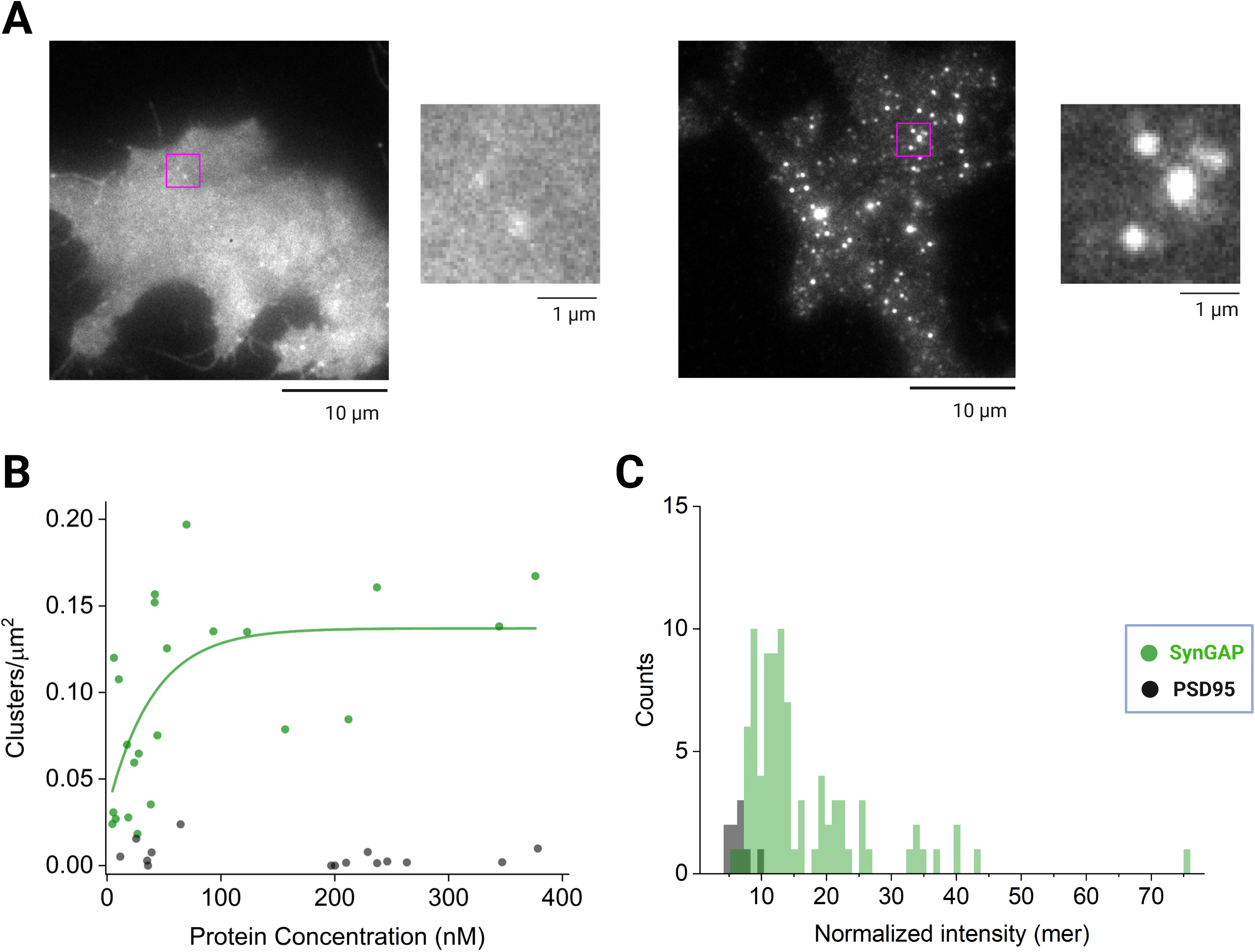
SynGAP forms nano-scale clusters in cells at nanomolar concentrations. (A) Representative cell images of PSD95-Halo (left) and Halo-SynGAP (right) in L cells (≈50 nM in the cytoplasm), with magnified images (of the area in magenta box) for each, showing typical clusters found at these concentration ranges. (B) Formation of nanoscale clusters of Halo-SynGAP (green) and PSD95-Halo (black) in L cells, examined as the clusters detected/µm^2^, as a function of total cytoplasmic protein concentration in each cell. Asymptotic fitting was performed to facilitate visualization and to estimate the concentration at which cluster density begins to plateau. The numbers of examined cells: 23 for SynGAP and 15 for PSD95. (C) The intensity distributions of all clusters detected in L cells expressing Halo-SynGAP (green) and PSD95-Halo (black) (5 cells each) in the concentration range 10 – 40 nM. Clusters consisting of ≥4 molecules were detected. The average intensity of a cytoplasmic monomer-reference molecule (raft-targeting motif of Lyn linked to Halo) was used to normalize the intensities.

In cells with an average cytoplasmic concentration of 27 nM, SynGAP forms clusters containing 5 – 40 molecules, with a median copy number of 13.1 (mean = 16.6 ± 1.1; n = 5 cells and 88 clusters). PSD95, at an average cytoplasmic concentration of 29.4 nM, hardly forms clusters, but the small numbers of clusters found under these conditions exhibited a median size of 6.7 molecules (mean = 6.6 ± 0.6; n = 5 cells and 9 clusters) (**Figure 4C**).

### SynGAP condensation is moderately suppressed by CaMKII

After LTP stimulation, CaMKII is activated and phosphorylates SynGAP, which might lead to the removal of SynGAP from post-synaptic sites (Araki et al., 2015). SynGAP has several known CaMKII phosphorylation sites, and interestingly, most are located in the IDR (Walkup et al., 2015). Therefore, we tested the effect of CaMKII on SynGAP condensation (**Figure 5A**). The co-expression of kinase-dead CaMKII (K42M; Yamagata et al., 2009) hardly affected SynGAP condensation. However, when the constitutively-active CaMKII (T286D; Giese et al., 1998) was co-expressed, SynGAP condensation was moderately suppressed. This was reproduced by SynGAP’s phosphomimetic mutant (SynGAPα1 with S D mutations of all the reported phosphorylation sites; Walkup et al., 2016; Araki et al., 2015). This moderate reduction of SynGAP’s condensate forming capability might contribute to the decreased AMPAR density in neuronal synapses, by its diffusion out of spines (Araki et al., 2015).

**Figure 5.**
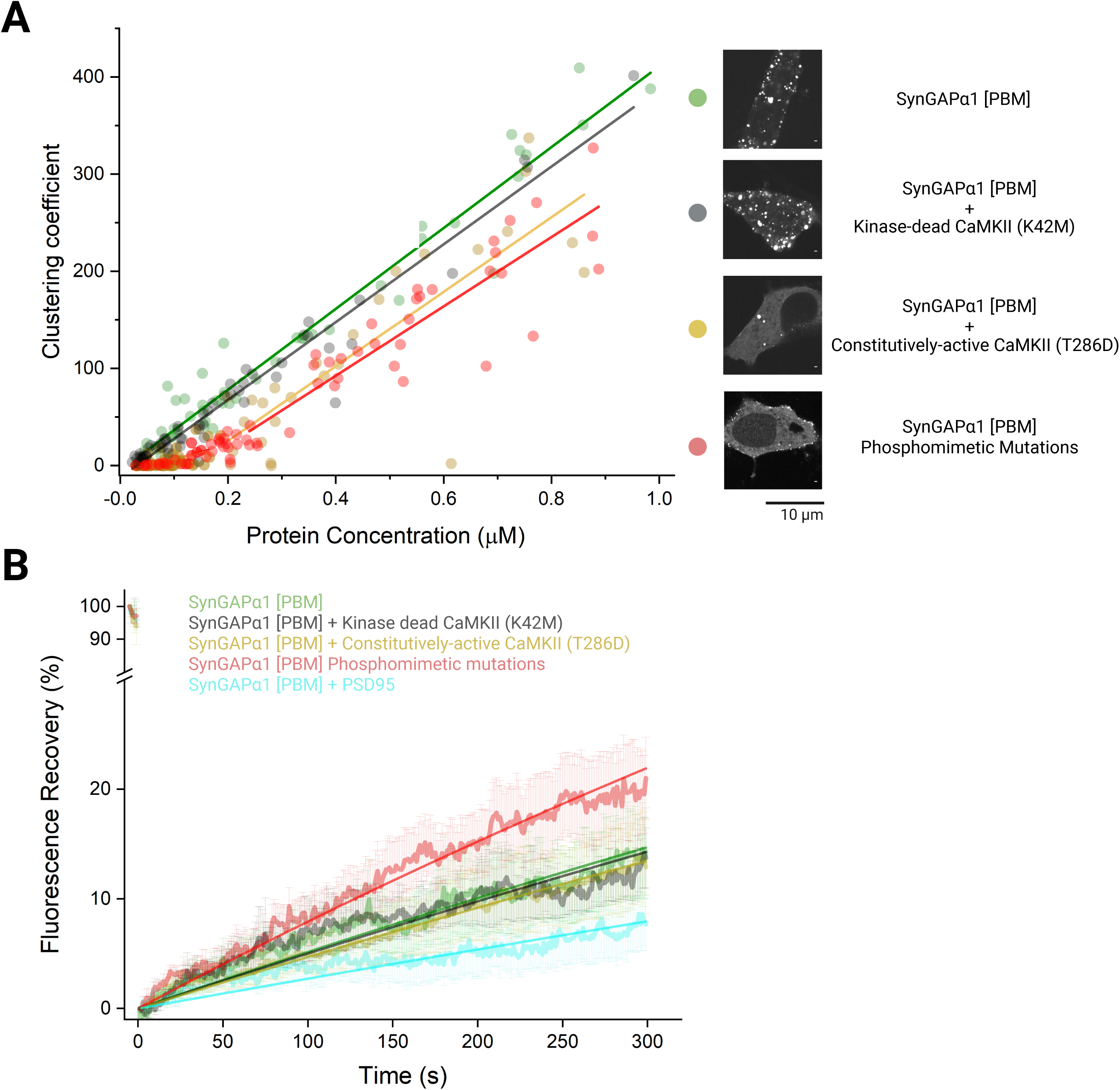
SynGAP condensation is moderately suppressed by CaMKII. (A) The effect of co-expressed CaMKII on the condensate formation of mGFP-SynGAPα1 (full length) in L cells (control, same data as those shown in Figure 3A; 47 cells) kinase-dead CaMKII, (57 cells); constitutively active CaMKII, (81 cells), and the condensate formation of the phosphomimetic mGFP-SynGAPα1 mutant (68 cells), displayed as a function of the protein concentrations in the cytoplasm. Linear regression was performed in the entire range for control and kinase-dead CaMKII expression and in the range of 0.3-1.1 µM for others. (B) The presence of PSD95 in SynGAP condensates decreases the liquidity of condensates, while phosphomimetic mutations increase it. FRAP recovery curves for the condensates of mGFP-SynGAP and its phosphomimetic mutants in L cells (≈1 µM in the cytoplasm), as well as for the condensates of mGFP-SynGAP in the presence of a kinase-dead or constitutively active mutant of CaMKII or PSD95.

SynGAP phosphorylation might also change the liquidity of its condensates. We tested this possibility using FRAP (**Figure 5B**). The condensates of SynGAP’s phosphomimetic mutant exhibited faster recovery (τ_1/2_ = 14 min) compared with those of wild-type SynGAP (τ_1/2_ = 21 min). However, the recoveries of the wild-type SynGAP condensates formed in the presence of the constitutively-active CaMKII or kinase-dead CaMKII (τ_1/2_ = 23.9 min and τ_1/2_ = 22.4 min, respectively) were essentially the same as that in the absence of CaMKII (τ_1/2_ = 21 min). Possibly, the number of residues phosphorylated by co-expressed CaMKII might be lower than the number of artificial S D mutations in the phosphomimetic mutant. The lower level of phosphorylation might still be sufficient to reduce the efficiency of condensate formation compared with that of wild-type SynGAP, but not able to increase the condensate liquidity to the level found in the condensates of the phosphomimetic mutant. Interestingly, co-expression of PSD95 with SynGAP at approximately equimolar concentrations caused slower FRAP (τ_1/2_ = 41.7 min), indicating that PSD95-bound SynGAP might undergo further gelation compared to SynGAP condensates without PSD95.

### SynGAP[PSD95] condensates immobilize Nlg1 in oligomerization-dependent manner

We investigated whether the SynGAP condensates formed in the presence of approximately equimolar concentrations of PSD95, termed SynGAP[PSD95] condensates in this report, recruit post-synaptic transmembrane receptors that bind to PSD95, mimicking the behavior of the PSD. First, we examined the recruitment of Neuroligin1 (Nlg1), a postsynaptic adhesion protein that directly binds to PSD95 via its C-terminal PDZ binding motif (Irie et al., 1997), to the condensates of full-length SynGAP[PSD95] formed in L cells. We found a Halo7-tag insertion position that did not affect the Nlg1 adhesion function, by using L cells expressing Halo7-conjugated Nlg1 and those expressing neurexin, the Nlg1 binding partner in the presynapse (**Supplementary Figures 4A and 4B**). The Halo7-tagged Nlg1 was fluorescently labeled and observed at the level of single molecules in live L cells containing SynGAP[PSD95] (average cytoplasmic concentration of ≈1 µM). Since wild-type Nlg1 (WT-Nlg1) was previously proposed to form a constitutive dimer (Arac et al., 2007), and the monomeric Nlg1 mutant (mNlg1; Dean et al., 2003; Arac et al., 2007) does not replicate the WT-Nlg1 function (Shipman et al., 2012), we also examined whether mNlg1 (**Figure 6A**) is recruited to SynGAP[PSD95] condensates. In addition, since Nlg1 oligomerization is required for its function to induce β-neurexin clustering to trigger the recruitment of synaptic vesicles, and perhaps to align the pre- and post-synaptic functional nano-domains (Dean et al., 2003; Giannone et al., 2013; Haas et al., 2018; Nozawa et al., 2022), we examined the recruitment of antibody-crosslinked mNlg1 (**Figure 6A**). The fractions of monomers, dimers, trimers, and tetramers, in terms of the numbers of molecules and fluorescent spots, were evaluated from the signal intensity distributions of individual spots (≈0.6 spots/µm^2^) and are summarized in **Figure 6A** (see **Supplementary Figure 4C**).

**Figure 6.**
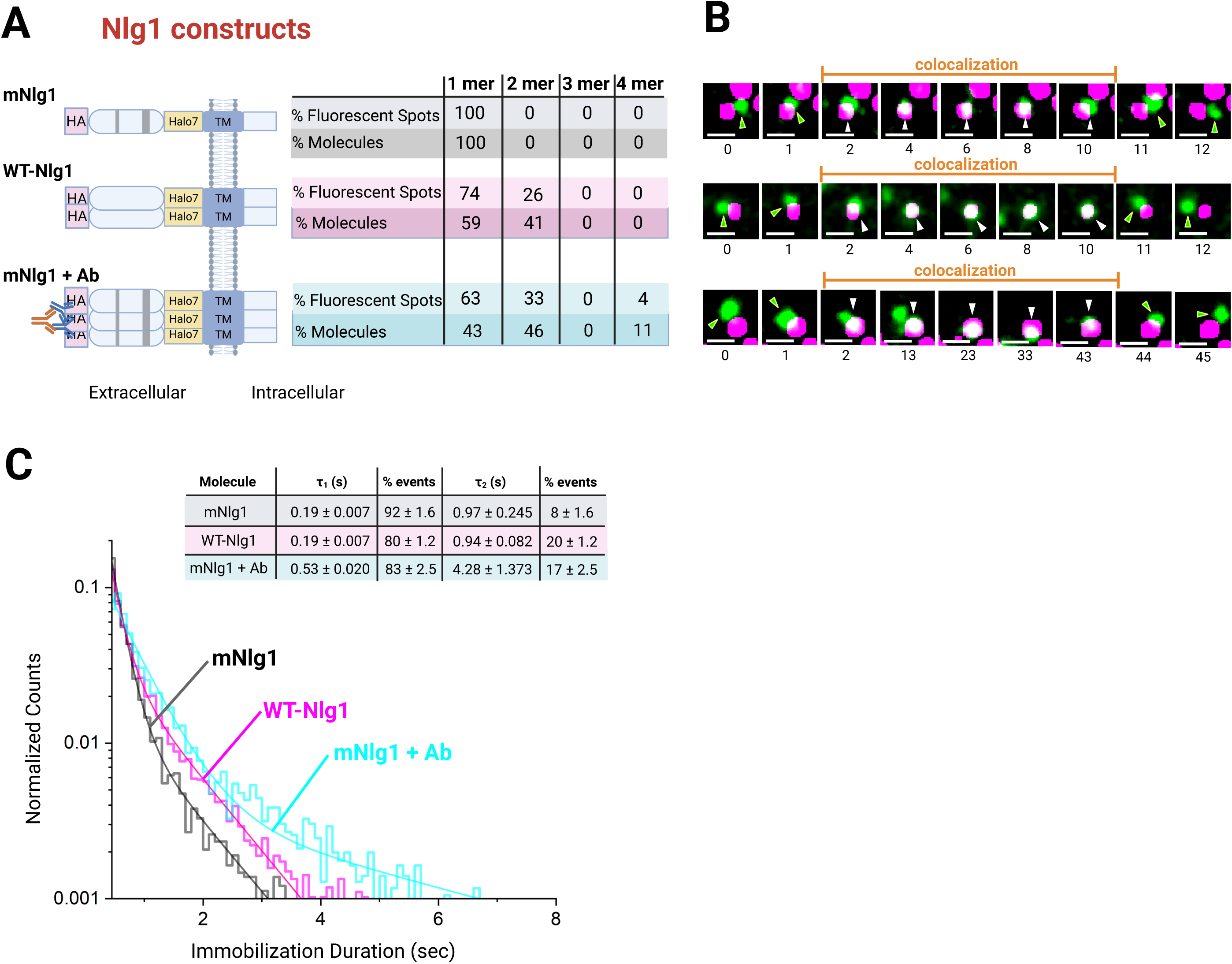
Nlg1 dynamically interacts with SynGAP[PSD95] condensates in an oligomerization-dependent manner. (A) Schematic figures of Halo-tagged mNlg1, WT-Nlg1, and mNlg1 crosslinked with primary and secondary antibodies (for the test of Halo-tagged WT-Nlg1 function, see **Supplementary Figure 4A**) with a table summarizing the fluorescent spot fractions and molecular fractions for Nlg1 monomers, dimers, trimers, and tetramers in the L-cell PM, expressing Nlg1s at a density of ≈0.6 spots/µm^2^. (B) Typical simultaneous two-color single-molecule image sequences, demonstrating that the Nlg1 molecules shown in A (green) become colocalized with and immobilized at SynGAP[PSD95] condensates (magenta). White arrowheads, colocalized and immobilized molecules; green arrowheads, diffusing molecules near the immobilization site. The numbers below the individual still images indicate the frame number in the image sequence (10 Hz observations). (C) The duration distributions of colocalization/immobilization of Nlg1 fluorescent spots at the SynGAP[PSD95] condensates. The distribution was operationally fitted by the sum of two exponential decay functions, providing short (t_1_) and long (t_2_) immobilization lifetimes and their number fractions in terms of the immobilization event (summarized in the inset table). For the statistical analysis results and the number of experiments, see Supplementary Tables 1 and 3.

The recruitment of these Nlg1 molecules to SynGAP[PSD95] condensates was observed in L cells at the level of individual fluorescent spots (**Figure 6B**). When they were recruited to the SynGAP[PSD95] condensates, their fluorescent spots became virtually immobile. Therefore, the dwell lifetimes of individual spots on the SynGAP[PSD95] condensates were evaluated by measuring the immobilization lifetimes after they became colocalized with the SynGAP[PSD95] condensates. The distributions of the residency durations of individual fluorescent spots at the SynGAP[PSD95] condensates (**Figure 6C**) were operationally fitted with the sum of two exponential decay functions, providing the shorter (τ_1_) and longer (τ_2_) dwell lifetimes and their molecular fractions (**Figure 6C**). Both the τ_1_ and τ_2_ dwell lifetimes of mNlg1 and WT-Nlg1 are about the same (even monomeric mNlg1 exhibited two dwell lifetime components), but the fraction of the τ_2_ dwell-lifetime component of WT-Nlg1 was 2.5x greater than that of mNlg1 (**Figure 6C**), which might be because the dimers tend to have greater τ_2_ component fractions (**Figure 6A, right**). The mNlg1+Ab spots exhibited much longer τ_1_ and τ_2_ values. The approximately 3x elongated τ_1_ value might be due to the unresolved τ_1_ and τ_2_ components found in WT-Nlg1, and the 4.5x elongated τ_2_ value for the mNlg1+Ab spots suggests that tetramers stay at SynGAP[PSD95] condensates much longer than monomers and dimers. (The analysis results of the histograms and the statistical test results are summarized in **Supplementary Table 1 and 3**.) These results indicate that SynGAP[PSD95] condensates dynamically concentrate Nlg1, with tetramers concentrated much more efficiently than monomers and dimers. This result is consistent with the previous finding that suggested Nlg1 oligomerization is required for its function (Dean et al., 2003; Giannone et al., 2013). Taken together, we conclude that Nlg1 assembly in the PSD is enhanced by both Nlg1 oligomerization and binding to SynGAP[PSD95] condensates. The dwell lifetimes observed here would be shorter than those in the synapse probably due to the lack of interaction with neurexin, which might induce further clustering (see the next subsection).

### SynGAP[PSD95] condensates also immobilize TARP2 in AMPA-induced oligomerization-dependent manner

Next, we studied whether the SynGAP[PSD95] condensates recruit the complex of the AMPA receptor (AMPAR) and trans-membrane AMPA receptor regulatory protein 2 (TARP2) (Shanks et al., 2010) in similar manners to those we observed for Nlg1 recruitment. The numbers and subunit compositions of AMPARs at the post-synapse are critically regulated for synaptic plasticity and sustaining basal synaptic activity (Shi et al., 2001; Lu et al., 2009; Henley et al., 2016), and AMPARs bind to PSD95 via TARP2, which, like Nlg1, contains a C-terminal PDZ binding motif. For this investigation, we employed the AMPAR subunit GluA1, which by itself can form AMPAR by tetramerization (Cull-Candy et al., 2006).

We examined TARP2, the N-terminal-domain-deletion mutant of GluA1 (capable of forming dimers; Morise et al., 2019) linked to TARP2 (GluA1(ΔNTD)-TARP2), and the wild-type GluA1 (capable of forming tetramers; Cull-Candy et al., 2006) linked to TARP2 (GluA1-TARP2), as shown in **Figure 7A**. In these constructs, TARP2 served as a binder to PSD95. TARP2 expressed in L cells was indeed found monomeric (**Figure 7A and Supplementary Figure 5A**) and as it diffused in the L cell PM, it was recruited to the SynGAP[PSD95] condensates (**Figure 7B**). The TARP2 dwell duration distribution on the SynGAP[PSD95] condensates was quite similar to that of WT-Nlg1 (**Figure 7C**). GluA1(ΔNTD)-TARP2 formed dimers and tetramers at levels comparable to mNlg1+Ab (compare **Figure 7A** with **Figure 6A**), but their dwell time distributions were similar to that of WT-Nlg1, which did not form oligomers greater than dimers. This might be due to lower binding affinity of TARP2 to PSD95, compared to that of Nlg1 to PSD95, consistent with previous reports (1.2 µM vs 0.23 µM; Irie et al., 1997; Vistrup-Parry et al., 2021). GluA1-TARP2 formed considerably more trimers and tetramers than GluA1(ΔNTD)-TARP2 and mNlg1+Ab (compare **Figure 7A** with **Figure 6A**), and exhibited extended τ_1_ and τ_2_ dwell lifetimes. However, these lifetime values and their fractions are similar to those found for mNlg1+Ab, which formed fewer trimers and tetramers than GluA1-TARP2, again suggesting that the binding affinity of TARP2 to PSD95 might be lower than that of Nlg1. Thus TARP2, like Nlg1, exhibits valency-dependent immobilization in SynGAP[PSD95] condensates. Taken together, these single molecule imaging studies of post-synaptic receptors suggest that receptor oligomerization may be an important mechanism in the regulation of receptor binding to PSD.

**Figure 7.**
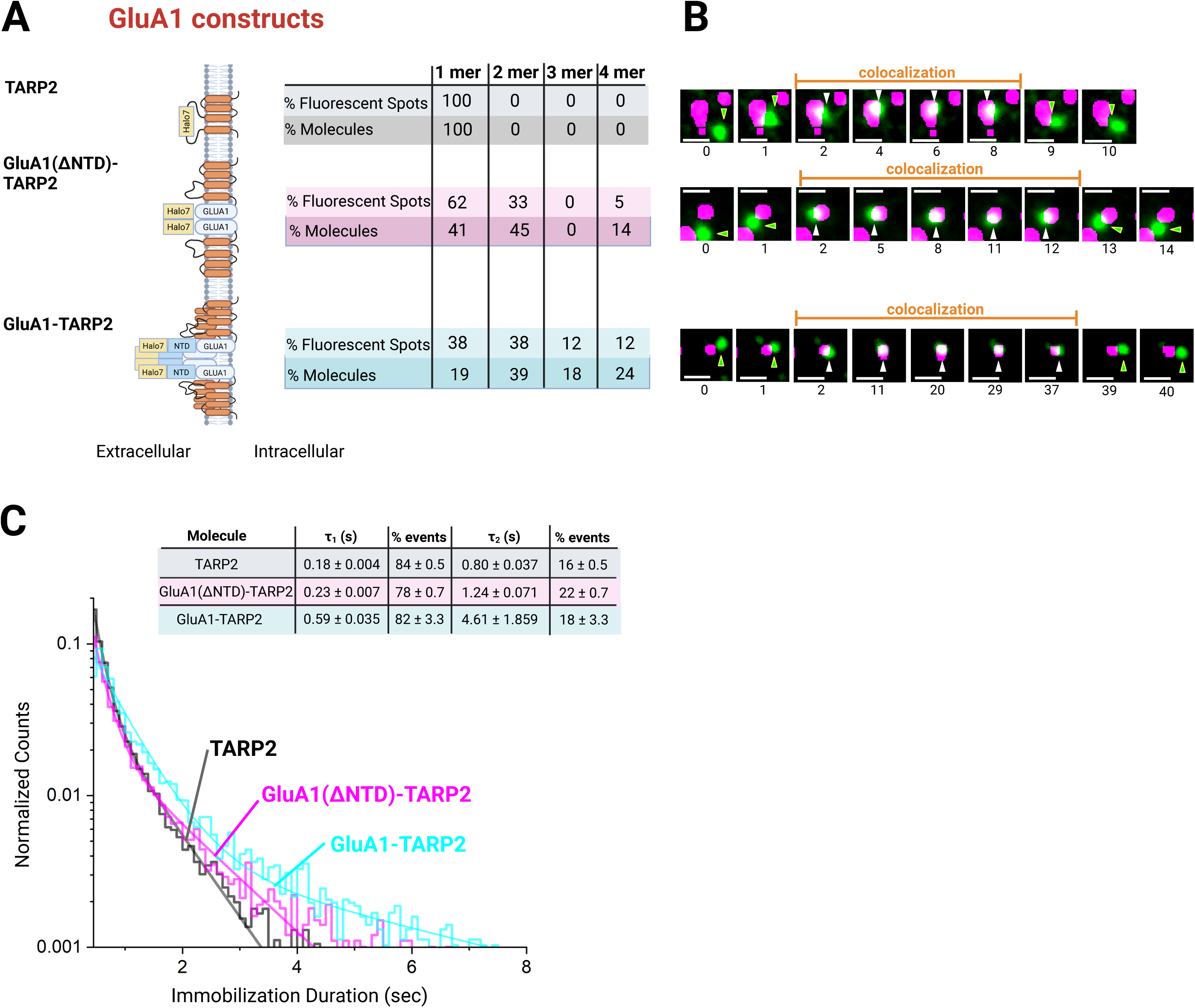
TARP2, GluA1(ΔNTD)-TARP2, and GluA1-TARP2 dynamically interact with SynGAP[PSD95] condensates in an oligomerization-dependent manner. (A) Schematic figures of Halo-tagged TARP2 and GluA1-linked TARP2, with a table summarizing the fluorescent spot fractions and molecular fractions for monomers, dimers, trimers, and tetramers of these constructs in the L-cell PM (at a density of ≈0.6 spots/µm^2^). (B) Typical simultaneous two-color single-molecule image sequences, demonstrating that the three TARP2 constructs shown in A (green) become colocalized with and immobilized at SynGAP[PSD95] condensates (magenta). White arrowheads, colocalized and immobilized molecules; green arrowheads, diffusing molecules near the immobilization site. The numbers below the individual still images indicate the frame number in the image sequence (10 Hz observations). (C) The duration distributions of colocalization/immobilization of the fluorescent spots of three TARP2 constructs at the SynGAP[PSD95] condensates. The distribution was operationally fitted by the sum of two exponential decay functions, providing short (t_1_) and long (t_2_) immobilization lifetimes and their number fractions in terms of the immobilization event (summarized in the inset table). For the statistical analysis results and the number of experiments, see **Supplementary Tables 2 and 4**.

### Dimerization of Nlg1 is required for proper synaptic localization, and Nlg1 can recruit SynGAP condensates to the PM and/or induce their formation on the PM

In the synapses of mature cultured rat hippocampal neurons (DIV16-22), the recruitment of mNlg1 to the post-synapse was much less efficient than that of WT-Nlg1 (**Figures 8A and 8B**). The difference in the recruitment between mNlg1 and WT-Nlg1 was much more evident in the neuronal synapse than expected from that in SynGAP[PSD95] condensates in L cells (**Figure 6C**). This is probably because, in the neuronal synapse, neurexin nanoclusters in the presynapse (Mondin et al., 2011; Chamma et al., 2016; Haas et al., 2018; Nozawa et al., 2022) would bind to Nlg1 dimers and induce Nlg1 clusters more efficiently compared to Nlg1 monomers, and the induced Nlg1 clusters in turn could be anchored to the PSD with enhanced efficiencies (assuming that mNlg1 does not form dimers with WT-Nlg1). Note that, in these experiments, the expression levels of tagged WT-Nlg1 and mNlg1 were much lower than the endogenous Nlg1 levels, which was made possible by the use of single-molecule imaging. These results are consistent with the previous observation that EPSC was not enhanced even after mNlg1 overexpression in hippocampal slice sections (Shipman et al., 2012).

**Figure 8.**
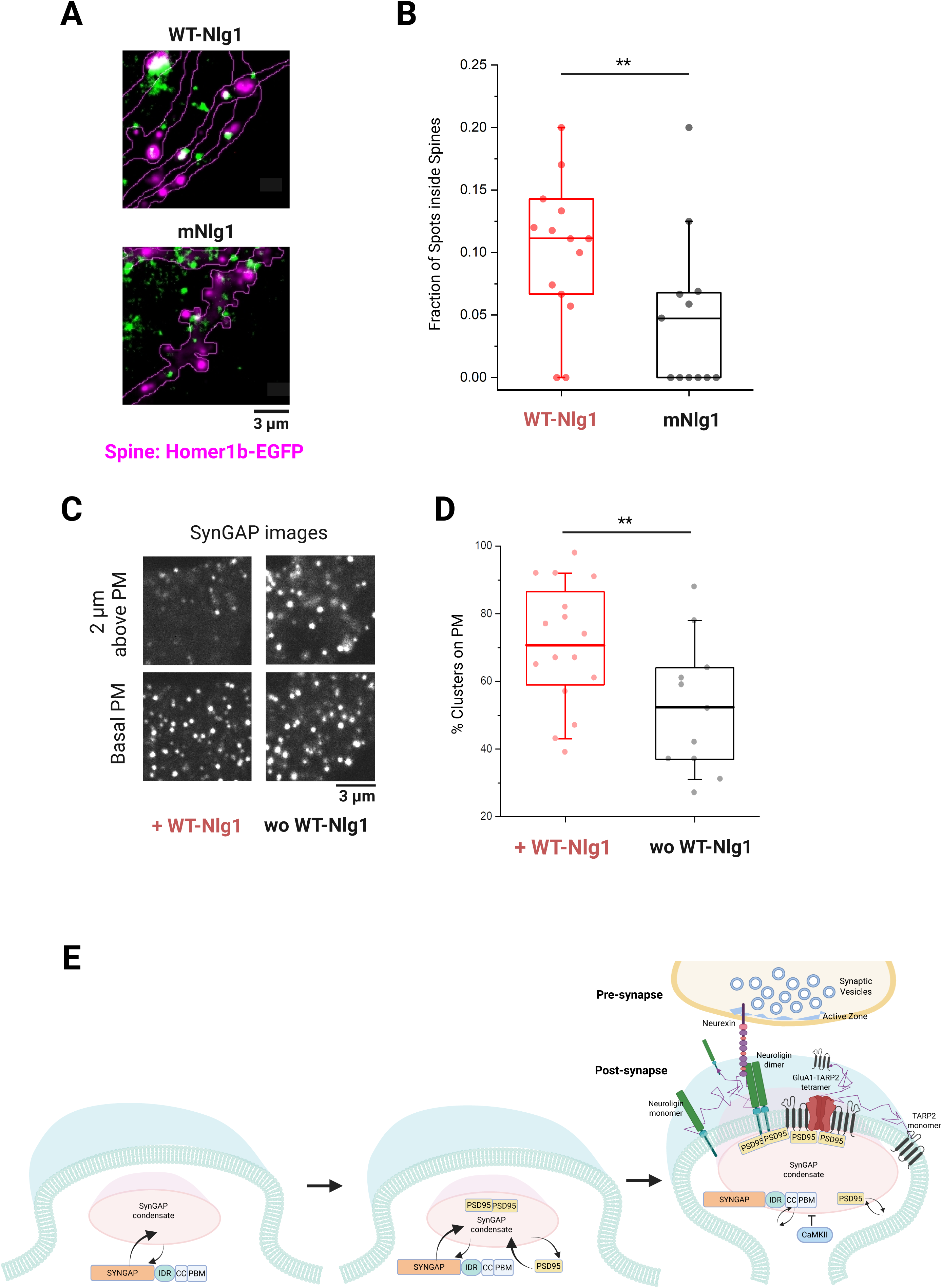
In neurons, Nlg1’s synaptic localization is enhanced by its dimerization, while the artificial SynGAP condensates in L cells are recruited to the PM by Nlg1 expression in the PM. (A) Representative TIRF images of the synaptic region marked with Homer1b-EGFP (magenta) and single molecules of WT-Nlg1 and mNlg1 (green) in mature cultured hippocampal neurons (DIV16-22). (B) Fluorescent spot fractions of single molecules of WT-Nlg1 and mNlg1 localized inside the synaptic regions marked by Homer1b-EGFP. Each dot represents the result obtained for a single neuron (5 independent replicates). Lateral bars, boxes, and whiskers indicate the mean values, inter-quartile ranges (25% - 75%), and 10% - 90% ranges, respectively, and asterisks indicate *p* < 0.05, using the two-tailed T-test (same in panel D). (C) Representative confocal microscopy images of SynGAP[PSD95] condensates in live L cells stably expressing WT-Nlg1 (left) and L cells without WT-Nlg1 expression (right), at the basal PM (bottom) and 2 µm above the basal PM (top). (D) Fractions of SynGAP[PSD95] condensates localized on the basal PM in live L cells stably expressing WT-Nlg1. Each dot represents the result obtained for a single cell (3 independent replicates). All condensates visible in the confocal images taken at the bottom PM and 2 µm above the PM were counted. (E) Schematic figure showing our proposed model for synapse induction involving SynGAP[PSD95] condensates. SynGAP starts forming condensates when its concentration exceeds 0.04 µM (≈5 µM in the mature spine), recruiting PSD95, which in turn triggers the interaction with Nlg. Nlg assembly might be further enhanced by the interaction with neurexin in the presynapse, leading to the nano-clustering of both neurexin and Nlg, which could enhance the growth/recruitment of SynGAP[PSD95] condensates, leading to the enhanced recruitment of TARP2 and AMPARs.

Furthermore, using L cells containing SynGAP[PSD95] condensates, we found that more condensates are localized on the PM inner surface in cells expressing Nlg1, compared to cells without Nlg1 expression (**Figures 8C and 8D**). This might be due to the recruitment of SynGAP[PSD95] condensates formed in the cytoplasm away from the PM and/or the formation of SynGAP[PSD95] condensates together with Nlg1 on the PM. The former type of condensate-membrane interaction has been extensively reported in the LLPS literature (Zeng et al., 2018; Zeng et al., 2019). In either case, in the neuronal post-synapse formation, Nlg1 recruitment to PSD and PSD recruitment of Nlg1 would occur cooperatively (**Figure 8E**). Similar processes might take place for AMPAR assembly in the post-synapse (**Figure 8E**).

## DISCUSSION

Our results show that SynGAP, a RasGAP protein that is highly enriched at the post-synapse, is capable of forming biomolecular condensates by itself, at low micromolar concentrations *in-vitro* and at sub-micromolar concentrations in live cells (**Figures 1 and 2**). SynGAP’s IDR trimerization induced by the CC domain is necessary and sufficient for its LLPS-induced condensation, and does not require the coexistence of PSD95 (**Figure 3A**). SynGAP condensates recruit PSD95 as a client by its binding to the SynGAP’s PBM (**Figure 2C**). These findings challenge the view that PSD95 is the main driver of PSD assemblies by LLPS (Zeng et al., 2016; Zeng et al., 2018; Zeng et al., 2019; Zhu et al., 2024), which might have arisen due to the use of only the SynGAP CC domain containing PBM, as a substitute for SynGAP. Our study highlights the importance of IDR in addition to the CC domain in SynGAP condensation, and thus emphasizes the necessity of including the IDR in the test sequences and using the full-length protein sequences to arrive at a more complete understanding of protein behaviors. Remarkably, SynGAP even forms nano-scale clusters, containing tens of molecules, at intracellular concentrations of tens of nanomolar (**Figure 4**), suggesting it may play a role in the initiation of nano-scale assemblies that have been observed at the post-synapse (MacGillavry et al., Nair et al., 2013; Hosy et al., 2017; Haas et al., 2018).

We performed the in-cell experiments quantitatively, by always measuring the intracellular SynGAP and PSD95 concentrations (**Figures 2, 3, 4, and 5**). To our knowledge, this level of quantification has never been done in LLPS work in cells, and our results underscore the importance of performing quantitative experiments for intracellular condensate formation.

Using single-molecule imaging and tracking, we found that SynGAP[PSD95] condensates formed in L cells can recruit Nlg1 and the TARP2-AMPAR complex diffusing in the PM in oligomerization-dependent manners (**Figures 6 and 7**). The oligomerization state of transmembrane proteins may be a key determinant in their proper localization at the post-synapse. This is particularly true for Nlg1, and perhaps for other transmembrane proteins in the postsynaptic PM that interact with presynaptic molecules that also form clusters (**Figure 8**). Together with previous results (Morise et al., 2019), which suggested that AMPAR forms metastable tetramers, we suggest that transmembrane proteins such as Nlg1 and AMPAR may be dynamically retained for longer durations as oligomers in the post-synapse, whereas their monomers and individual subunits can escape from the post-synapse more readily.

Moreover, SynGAP condensation and Nlg1 oligomerization might occur cooperatively: SynGAP[PSD95] condensation by LLPS on the PM could be enhanced by Nlg1 oligomerization, which might work as seeds for LLPS, and the SynGAP[PSD95] condensates on the PM in turn would recruit more Nlg1 oligomers, which will further enhance SynGAP[PSD95] condensation and/or the recruitment of SynGAP[PSD95] condensates formed in the cytoplasm away from the PM (**Figure 8E**). In neuronal synapse formation, presynaptic nanoclustering of neurexin would also greatly contribute to Nlg1 oligomerization and clustering. Previously, using cultured hippocampal neurons, Araki et al. (2015) found that, upon synaptic stimulation, the amounts of SynGAP in the PSD are decreased, and the entrance of AMPARs increased in the post-synapse. Our results showing the reduction of SynGAP condensates by CaMKII (**Figure 5A**) found in L cells are consistent with the initial decrease of SynGAP amounts in the PSD in neurons (Araki et al., 2015), but the later increase of the AMPAR density in the postsynaptic membrane cannot directly be explained. Perhaps, at later time points, SynGAP assembly might be recovered and further enhanced by the continued entry of AMPAR, and possibly by that of Nlg1, into the postsynaptic region.

These observations suggest that SynGAP’s abilities to undergo LLPS to form condensates and to bind to PSD95 play critical roles in the regulation of excitatory synapse functions. Furthermore, we propose that at the initial stages of synapse formation, SynGAP[PSD95] condensation may play a leading role to initiate postsynaptic assembly due to its ability to form nano-scale clusters at concentrations around 10 – 100 nM in the cell, as well as to the ability of condensates to cooperatively assemble Nlg1 together with the neurexin nano-assemblies in the presynapse.

## ACKNOWLEDGEMENT

We thank Dr. Yuri Nemoto and Ms. Hiroko Hijikata for their kind instructions for the primary culture of rat hippocampal neurons and Profs. Ikuko Koyama-Honda and Noboru Mizushima of the University of Tokyo for their kind help and instruction about the methods for producing viral vectors and viruses and cell infection. We thank Prof. Peter Scheiffele of University of Basel, Prof. Shigeo Okabe of the University of Tokyo, Prof. Tomoo Hirano of Kyoto University, and Prof. Michiyuki Matsuda of Kyoto University for their kind gifts of Nlg1 cDNA, PSD95 cDNA, pCAGplay vector, and pBpuro and pCMV-mPbase plasmids, respectively. We also thank the members of the Kusumi laboratory, Milovanovic laboratory, and Kennedy laboratory for helpful discussions. This work was supported in part by Japan Society for the Promotion of Science (JSPS) Grants-in-Aid for Scientific Research (Kiban A to A.K. [21H04772], Kiban S to A.K. [16H06386], Kiban B to T.K.F. [16H04775], Kiban C to R.S.K. [17K07333], Wakate to T.A.T. [21K15058], and Challenging Exploratory Research to T.K.F. [18K19001] and A.K. [22K19334]), and a Japan Science and Technology grant ACT-X to T.A.T. (JPMJAX211B). WPI-iCeMS of Kyoto University is supported by the World Premiere Research Center Initiative (WPI) of MEXT.

## AUTHOR CONTRIBUTIONS

S.A. and A.K. conceived and formulated the project. S.A., T.A.T., C.H., T.K.F., M.B.K., D.M., and A.K. designed the biological experiments and participated in discussions. S.A. performed virtually all of the microscopy and single-molecule imaging experiments and the data analysis. T.A.T performed the labelling efficiency experiments and data analysis. T.A.T. and T.K.F developed single-molecule analysis software. S.A., T.A.T., C.H., M.B.K., D.M., A.K. evaluated the data. S.A and I.M. produced virtually all the cDNAs. S.A and A.N. performed neuron culture. S.A., I.M., C.H., P.G.A., T.M., and W.G.W., produced and isolated SynGAP, PSD95, and their derivatives. S.A., M.B.K., D.M., and A.K. wrote the manuscript, and all authors discussed the results and participated in revising the manuscript.

## COMPETING INTERESTS

The authors declare no competing interests.

## MATERIALS AND METHODS

### Purification of proteins

mGFP-SynGAP IDR-CC [PBM] (called mGFP-SynGAP-ICP; from human SynGAP, a.a. 16-1308 of NCBI accession KAI2541928.1) and PSD95-mScarlet (rat full length PSD95, Uniprot ID P31016) were prepared as outlined in **Supplementary Figure 1A**. The SynGAP-ICP cDNA was linked with N-terminal 6xHis tag, maltose binding protein (MBP), Tobacco Etch Virus protease digestion sequence (TEV), and mEGFP tag cDNA sequences and this product was cloned into the pEGFP vector. For PSD95, instead of mEGFP, mScarlet was employed, and conjugated at the C-terminus of PSD95. These chimeric molecules were expressed in Expi293F cells (Thermo Fisher Scientific A14635) for three days, following the manufacturer’s recommendations. The cells were harvested at 25°C and lysed by three freeze-thaw (37°C) cycles in buffer A, containing 200 mM NaCl, 1 mM ethylenediamine tetraacetic acid (EDTA) and 0.5 mM Tris(2-carboxyethyl)phosphine (TCEP), buffered at pH 7.4 with 25 mM 2-[4-(2-Hydroxyethyl)-1-piperazinyl]ethanesulfonic acid (HEPES), and supplemented with protease inhibitor cocktail (cOmplete Mini, Roche 11873580001), 10 µg/mL DNase I (Thermo Scientific EN0521), and 1 mM MgCl_2_ (EDTA: Dojindo 345-01865, TCEP: Fujifilm 209-19861, and HEPES: Dojindo 346-08235).

The lysates were centrifuged at 30,000g at 4°C for 1 h, and then the mGFP-SynGAP-ICP and PSD95-mScarlet proteins in the supernatants were purified by one-step and two-step affinity column chromatography protocols, respectively (**Supplementary Figure 1B**). The first step for both proteins was affinity purification on an MBP-binding dextrin Sepharose column (MPBTrap, GE Healthcare 28-9187-78). The lysates were loaded twice on the column at a flow rate of 0.5 mL/min, washed with 2 mL of buffer A, and then eluted with 10 mM maltose in buffer A at 25°C. Due to the ready condensation of mGFP-SynGAP-ICP after MBP was cleaved by the TEV protease, and since the purity of 6xHis-MBP-TEV site-mGFP-SynGAP-ICP after the first column chromatography was reasonably high, we stopped the mGFP-SynGAP-ICP purification at this step, and the 6xHis-MBP-TEV site was removed with His-tagged TEV protease, prepared as previously described (Tropea et al., 2009), right before the fluorescence microscopy experiments.

For the PSD95-mScarlet isolation, after the same steps of lysate preparation and first MBPTrap column chromatography for the preparative isolation of mGFP-SynGAP-ICP, we conducted the second purification step for separating PSD95-mScarlet from 6xHis-MBP after the TEV protease cleavage of the eluate from the first column chromatography (1 His-tagged TEV:10 total protein in the eluate by weight; at 16°C for 18 h), using Ni-NTA chromatography (HisPur™ Ni-NTA Resin, Thermo Scientific 88222). Before column chromatography, NaCl and imidazole were added to the digested solution at final concentrations of 400 and 25 mM, respectively, and then the solution was passed twice over a pre-equilibrated Ni-NTA Sepharose slurry (200 mL of bed volume, equilibrated with buffer containing 300 mM NaCl, 25 mM imidazole, and 0.5 mM TCEP buffered with 25 mM tris(hydroxymethyl)aminomethane (Tris)-HCl at pH 7.4 (Tris; Wako 011-16381), in a Poly-Prep Chromatography Column (Bio-Rad 731-1550). The flow-through was collected.

The mGFP-SynGAP-ICP eluate from the first column chromatography and the flow-through containing PSD95-mScarlet from the second column chromatography were concentrated with a Pierce Protein Concentrator (30 kDa MWCO, PES-membrane, Thermo Scientific: 88502) to ∼100 mL. The concentrated solution of PSD95-mScarlet was dialyzed against buffer B (150 mM NaCl and 0.5 mM TCEP buffered with 25 mM Tris-HCl at pH 7.4) to lower the salt concentrations and remove the imidazole, using micro-dialyzer units (10 kDa MWCO, Slide-A-Lyzer MINI, Thermo Scientific 69576) at 4°C overnight. Finally, the protein concentrations were determined using the absorbance at 280 nm, assuming molecular extinction coefficients of 10.61 and 12.26 x 10^4^ (/M/cm) for PSD95-mScarlet and 6xHis-MBP-TEV site-mEGFP-SynGAP-ICP, respectively (determined from the amino acid composition data). The purities of SynGAP-ICP and PSD95 were 77 and 76%, respectively, as determined by staining the SDS-PAGE gels with InstantBlue Coomassie Protein Stain (ISB1L, Abcam ab119211). Aliquots (10 µL) of the protein solutions were placed in Eppendorf tubes, rapidly frozen in liquid N_2_, and then stored at -80°C until use.

### *In-vitro* protein reconstitution experiments

Purified 6xHis-MBP-TEV site-mEGFP-SynGAP-ICP in the frozen stock was thawed at 37°C, and then incubated with TEV protease (1:50 total protein weight) at room temperature for 30 minutes to remove the MBP prior to *in vitro* reconstitution experiments. This solution and the PSD95-mScarlet solution were diluted with buffer B to the desired protein concentrations. After adding the selected amounts of PEG8K in buffer B, the mixtures were transferred to glass-bottom slides (Ibidi 81507) and observed by fluorescence confocal microscopy within 2 min after the PEG addition. All of these reactions and observations were performed at 25°C.

### cDNA constructs

The cDNAs encoding mouse Nlg1 and Neurexin 1β (Nxn) were PCR amplified from the pCAG-NL1AB and pCAG-HA-Nrxn1β AS4(-) vectors, which were gifts from Prof. Peter Scheiffele of the University of Basel (Addgene plasmid 15262 http://n2t.net/addgene:15262; RRID:Addgene_15262; Addgene plasmid 59409 http://n2t.net/addgene:59409; RRID:Addgene_59409). The cDNA encoding PSD95 was a gift from Prof. Shigeo Okabe of the University of Tokyo, Graduate School of Medicine (Okabe et al., 1999). The cDNA encoding SynGAP was acquired from the PlasmID repository at Harvard Medical School.

The sequences encoding various proteins were subcloned into the following vectors: pEGFP (Clontech Z2370N), pCAGplay (a custom vector containing the CMV enhancer, chicken β-actin promotor, and rabbit β−globin polyadenylation signal region; a gift from Prof. Tomoo Hirano of Kyoto University) (Kawaguchi and Hirano, 2007), and pPBpuro (a custom vector with the PiggyBac terminal repeat sequence and the high expression promoter consisting of chicken β-actin promoter, CAG promoter with an artificial intron, and CMV IE; a gift from Prof. Michiyuki Matsuda of Kyoto University) (Nakamura et al., 2020).

### Culture of L cells and HeLa cells and transfections

L cells and HeLa cells were cultured in Dulbecco’s Minimum Essential Medium (DMEM), purchased from Sigma Aldrich (D5796) and supplemented with 10% (vol/vol) fetal bovine serum (FBS, Gibco: 10270-106, Lot#: 42G7281K). All cells were mycoplasma-free, as determined by using MycoAlert (Lonza LT07-118) and a Mycoplasma PCR Detection Kit (Beyotime Biotechnology C0301S). The cells were transfected with the plasmid vectors by lipofection using the Effectene reagent (Qiagen 301425) with approximately 1 µg of cDNA encoding each protein, according to the manufacturer’s recommendations.

### Preparation of cell lines stably expressing proteins

For the generation of cell lines stably expressing selected proteins, the cells were transfected with the pPBpuro expression vector, and two days after transfection, the cells were incubated with medium containing puromycin (2 µg/mL) on 10 cm cell-culture grade dishes. After an incubation in the selection medium for a week, cell colonies were partly detached with 0.25% trypsin and picked up using a pipette tip, and the cells from these colonies were replated on glass-bottom dishes (described later). The expression levels of isolated cell clones were examined by fluorescence microscopy.

### Neuron culture and viral transfections

Primary hippocampal neurons were essentially cultured as described (Nakada et al., 2003) with slight modifications. Briefly, the glass-bottom dishes were coated overnight with 0.01% (w/v) poly-L-lysine (Sigma-Aldrich P2636) dissolved in water, and subsequently washed with doubly distilled water five times. A feeder layer of astrocytes isolated from rat hippocampi (Kaech et al., 2006) was plated on the circular plastic part of the glass-bottom dishes in DMEM containing 15% FBS, the day before starting the neuron culture. Hippocampi from E17-E19 Wister rat embryos were dissected and dissociated following the protocol described in the Pierce Primary Neuron Isolation Kit (Thermo Fischer MAN0016221). The isolated primary neurons were plated at a density of approximately 4 x 10^5^ cells/dish on the coverslip part of the glass-bottom dish in 2 mL MEM, containing 2% (v/v) B27 (Gibco 17504-044) and 5% (v/v) FBS. At 24 h after the neurons were plated, 1-β-D-arabinofuranosylcytosine (Ara-C; Sigma-Aldrich: C6645-25MG) was added to a final concentration of 1 µM, to prevent the proliferation of glial cells. These neurons were healthy and survived up to 6 weeks, and neurons between DIV 18 - DIV 21 were employed for our experiments. The microscope observations were made after the culture medium was replaced with artificial cerebrospinal fluid (ACSF; 146 mM NaCl, 2.5 mM CaCl_2_, 3 mM KCl, 1 mM MgCl_2_, 17 mM glucose) buffered with 10 mM HEPES (Gibco 15630-080) at pH 7.4.

Target proteins were expressed in primary hippocampal neurons by lentivirus infection. The lentivirus particles were produced by transfecting LentiX 293T cells (Takara 632180) with the pLVSIN-CMV Pur vector (Takara 6183) containing the cDNAs encoding the respective target proteins together with the Lentiviral High Titer packaging mix (Takara 6194), and then isolated according to the manufacturer’s instructions. Lentivirus particles were added to primary hippocampal neurons at DIV 6-7.

### Antibody-induced clustering of mNlg1

Clusters of mNlg1 were induced by adding a combination of primary antibodies against its N-terminal HA tag (anti-HA monoclonal antibody, Cell Signaling Technology C29F4, 1:1,000 dilution) and Halo tag (anti-HaloTag polyclonal antibody, Promega G9281, 1:500 dilution) into the imaging medium, 30 minutes before observation.

### Fluorescence labeling of Halo-tagged proteins

For the TMR fluorescence labeling of Halo-tag proteins fused to target proteins, the cells were incubated in complete growth medium containing 20 nM TMR-Halo linker (Promega G8251) at 37°C for 30 min. After labeling, cells were washed four times with imaging medium prior to imaging. When Halo-tagged Nlg1 expressed in primary hippocampal neurons was labeled with TMR, the neuron culture medium containing 20 nM TMR-Halo linker was used for the labeling, and the cells were then washed six times with ACSF buffered with HEPES at pH 7.4 prior to imaging.

The labeling efficiency with TMR was estimated in the following way. Plasma membrane signal of Halo-tagged CD47 protein, in live cells, was measured with increasing incubation periods with TMR dye. In the observation period after 60 min, the concentration of TMR dye was doubled, to ensure that no unlabeled protein remained. Incubation with TMR dye at a concentration of 5 nM or higher, for a period of 30 min, was sufficient to label almost all CD47 molecules. To estimate the percentage of Halo and mGFP tagged molecules that were actually fluorescent, taking into account misfolding, slow folding, and cleavage of the tagged proteins, a monomeric molecule, with both Halo and mGFP tags, GFP-CD47-Halo (intracellular Halo tag) was labelled with the above saturating condition, and the percentage of Halo spots colocalized with mGFP spots, and the percentage of mGFP spots colocalized with Halo spots was estimated using single-molecule imaging and colocalization analysis. The percentages of fluorescent proteins were estimated to be 88 ± 2.2% for Halo7, and 76 ± 2.3% for mEGFP. Labeling efficiencies of TMR labeling for transmembrane proteins as well as for Halo-SynGAP were performed to achieve maximum labelling efficiency for experiments to estimate fractions of monomers, dimers, and oligomers.

In the single-molecule tracking experiments, transmembrane proteins were labeled with SeTau647 so they could be observed together with the condensates of mCherry-SynGAP and PSD95-mGFP. For SeTau647 labeling, the cells were incubated with 60 nM SeTau647-Halo linker (custom ordered from Shinsei Kagaku) at 37°C for 30 min. After labeling, the cells were washed four times with imaging medium prior to imaging.

### Confocal fluorescence microscope imaging

For fluorescence confocal microscopy, the L cells and HeLa cells were plated in glass-bottom dishes with a 12 mm diameter glass window (No.1S glass, Iwaki 3971-035), pre-coated with fibronectin (FN, Sigma Aldrich F1141) by placing 100 µL of 10 μg/mL FN in phosphate buffered saline (PBS: 137 mM NaCl, 8.1 mM NaHPO_4_, 2.7 mM KCl, 1.5 mM KH_2_PO_4_, pH 7.4) on the glass at room temperature for 10 min. The cells were cultured in complete growth medium for 24–48 h before imaging. For microscope observations, the medium was replaced with a specially designed Ham’s F12 medium, lacking phenol red, NaHCO_3_, and glutamine, but containing 1 mM GlutaMAX and lower concentrations of the following ingredients: 10 nM Cu(II)SO_4_, 30 nM D-biotin, and 300 nM linoleic acid. The medium was buffered with 2 mM TES (Dojindo 348-08371) at pH 7.4 and supplemented with a final concentration of 10% (v/v) filtered FBS (imaging medium).

For observations of primary neurons in culture, the culture medium was replaced with ACSF buffered with 10 mM HEPES at pH 7.4.

To obtain confocal fluorescence images of the cells at 37°C, we used an Olympus IX83 inverted microscope equipped with a Yokogawa spinning disk confocal unit, CSU-W1, an Olympus objective UApoN100XTIR, and a Hamamatsu Photonics ORCA-Flash 4.0 V2 plus sCMOS camera. Confocal image slices were taken every 0.5 µm, starting from the basal plasma membrane. The entire microscope was housed within a home-built environmental chamber (Tsunoyama et al., 2018).

### Evaluation of protein clustering using “clustering coefficient”

To evaluate protein clustering, confocal slice images were taken every 0.5 µm for a total of 4 slices from right above the basal plasma membrane of each cell, and then analyzed to detect clusters and measure the total cell area (excluding the nucleus) using the open-source machine-learning based imaging software, ‘ilastik’ (Berg et al., 2019). The “clustering coefficient” was defined as the total intensity of all pixels identified as ‘clusters’ in an image, divided by the total area of the cell in pixels. This value should be proportional to the expected protein concentration in the cytoplasm if the protein in the condensates were dispersed throughout it. Under the phase separation conditions, as the protein concentration is increased, the protein concentrations in the condensates and in the bulk cytoplasm will remain the same, and therefore, the *amount* of the protein in the condensate will linearly increase with increasing protein concentration, after the protein concentration in the cell reaches the level at which phase separation starts occurring. Therefore, the clustering coefficient will also increase linearly with increasing protein concentration. In the present study, the clustering coefficient is plotted as a function of the average concentration of the protein (pixel intensity).

### Evaluating the mGFP and Halo bound SynGAP and PSD95 concentrations in the cytoplasm by confocal microscopy

To quantitatively evaluate the concentrations of mGFP-bound SynGAP-ICP and PSD95 in the cytoplasm of L cells by confocal microscopy, the concentration calibration curves were obtained in the following way. A known amount of purified EGFP (Biovision 4999) was reconstituted in imaging buffer, and EGFP solutions covering the concentration ranges in L cells were produced by dilution. These EGFP solutions were placed in the glass-base dishes used for observing the cells, and imaged with the same confocal microscope and the same instrument settings employed to observe L cells for evaluating the clustering coefficient. Three totally independent determinations were made. The average pixel intensity of each EGFP solution was plotted as a function of the protein concentration (**Supplementary Figure 2A**). The data set was fitted with a linear function, which was then used to convert the average pixel intensities of mEGFP-bound SynGAP-ICP and PDS95 in the cytoplasm to the actual concentrations.

The concentrations of Halo-bound SynGAP and PSD95 in the cytoplasm of L cells were evaluated in a similar way as for mGFP-bound proteins. Purified Halo protein (Promega G4491) was labelled with TMR-Halo linker at 5-fold molar excess overnight at 4°C, and the TMR labelled Halo protein was separated from the unreacted dye by passing it through a PD10 desalting column (Cytiva 17085101). The amount of TMR-labelled Halo protein was evaluated by absorbance measurements at 556 nm using the extinction coefficients for TMR-labelled Halo protein previously determined (Liu et al., 2024). The labelled Halo protein was imaged as described above for EGFP protein, and average pixel intensity for each Halo protein was plotted as a function of the protein concentration (**Supplementary Figure 3A**).

### Single-molecule fluorescence imaging and analysis

All single-molecule observations were performed at 37°C, using a home-built objective-lens-type TIRF microscope housed in home-made environmental chambers, as described previously (Koyama-Honda et al., 2005, 2020; Tsunoyama et al., 2018). Briefly, the TIRF microscope was built on an Olympus inverted microscope (IX-83) equipped with a 100x 1.49 NA objective lens (UApoN100XTIRF). The three-color fluorescence images were separated into the three detection arms of the microscope by two dichroic mirrors (Chroma ZT 561rdc-UF3 and ZT640rdc-UF3). The detection arms were equipped with bandpass filters of ET525/50m, ET600/50m, or ET700/75m (all from Chroma), and the images were recorded at each arm with a two-stage microchannel plate intensifier (Hamamatsu Photonics C9016-02MLP24), lens-coupled to a scientific complementary metal oxide semiconductor (sCMOS) camera (Hamamatsu Photonics ORCA-Flash ver4.0 V2 plus C11440-22CU). The final magnification was 133x, resulting in a pixel size of 48.75 nm for all channels. Only the 642 nm channel was used for single-molecule observations, while the 488 nm and 561 nm channels were used to image PSD95-EGFP and mCherry-SynGAP clusters, respectively. The incident excitation laser powers at the sample plane were 0.07 µW/µm^2^ for the 488 nm line (for mEGFP, Coherent OBIS 488-100 LS), 0.06 µW/µm^2^ for the 561 nm line (for TMR, Coherent OBIS 561-100 LS), and 0.4-0.5 µW/µm^2^ for the 642 nm line (for single molecules of SeTau647, Omicron LuuXPlus 640-140). For triple-color experiments, the 488 nm and 561 nm channels were used for imaging PSD95-EGFP and mCherry-SynGAP clusters, respectively, at a frame rate of 1 Hz with an exposure time of 33 ms, whereas the 642 nm channel was used for imaging single molecules of Halo-tagged transmembrane proteins at a frame rate of 10 Hz with an exposure time of 33 ms. All three channels were staggered in time with 33-ms intervals between successive images of each channel, to minimize cross channel fluorescence signals.

Each individual fluorescent spot in the image was identified and tracked by using an in-house computer program, developed by Fujiwara et al. (2002; 2023). The superimposition of images in different colors obtained by three separate cameras was conducted using the method developed by Koyama-Honda et al. (2005). Transient immobilization events were detected in single-molecule trajectories by using the same program as for single fluorescent spot tracking, and by setting the upper threshold of the diffusion coefficient defined for a single step (0.1 s) to 0.0083 µm^2^/s and the lower threshold of the number of frames to five (0.5 s).

To detect immobilizations of single fluorescence spots of transmembrane proteins within SynGAP condensates, SynGAP images obtained using the 561 nm channel were binarized using a custom-made ImageJ script. The binarized SynGAP channel masks were then used in an in-house computer program to detect whether the trajectories of immobilized spots were located inside or outside the binarized areas. Histograms of immobilization durations for each transmembrane protein were fitted with the sum of two exponential decay functions in the range from 0.4 s to 200 s.

The fractions of monomers, dimers, trimers, and tetramers of transmembrane proteins were obtained following the method described previously (Morise et al., 2019). First, the distribution of the signal intensities of monomeric molecules was obtained by employing the Halo-tagged transmembrane domain of LDL receptor (Halo7-TM) labeled with SeTau647. The signal intensities of all the individual fluorescent spots detected in the initial twenty frames in single fluorescent-molecule imaging movies were determined, to obtain the signal intensity histogram. The histogram could be fitted well using a single lognormal function. Then, the signal intensity distributions of the fluorescent spots of transmembrane proteins of interest were obtained and fitted by the sum of four lognormal functions, based on the mode and standard deviation of the lognormal function for the monomer signal intensity distribution. Oligomers greater than tetramers were not detected.

Under these expression conditions, we found that only 41% of WT-Nlg1 molecules exist as dimers, while 59% of molecules (protomers) exist as monomers, at variance with the prevalent notion that all Nlg1 molecules constitutively exist as homodimers. Perhaps, the high dimer content would be true where the number densities of Nlg1 are very high, such as in the PSD. Under similar expression conditions, the mNlg1 mutant mostly existed as monomers, whereas antibody-crosslinked mNlg1 (mNlg1 + Ab) exhibited more dimers and tetramers compared with the wild-type Nlg1 (**Supplementary Figure 4B** and **Figure 4A**). For the GluA1(ΔNTD)-TARP2 molecules, 41%, 45%, and 14% existed as monomers, dimers, and tetramers, respectively, while 19%, 39%, 18%, and 2% of GluA1-TARP2 molecules existed as monomers, dimers, trimers, and tetramers, respectively.

### Single-molecule fluorescence imaging and analysis for nanoscale clusters

To detect nanoscale clusters of Halo-SynGAP (and PSD95-Halo) labelled with TMR in L cells, fluorescence spots on the plasma membrane were imaged as described above, in the 561nm channel, at laser intensities 1x, 2x, and 4x lower than those used for single-molecule detections of the monomer molecule (raft-targeting motif of Lyn linked to Halo). The signal intensities of all the individual fluorescent spots detected in the initial ten frames in single fluorescent-molecule imaging movies were determined, to obtain the signal intensity histogram. The signal intensities were normalized by the mean intensity obtained from histogram of monomer molecules.

### Fluorescence recovery after photobleaching (FRAP)

For the in-cell fluorescence recovery after photobleaching (FRAP) experiments, a home-built TIRF microscope based on a Nikon inverted microscope (TiE), equipped with a focused 488-nm laser system for photobleaching (Coherent OBIS 488-100 LS, Maximum power: 150 mW) and 100x 1.49 NA objective lens (ApoTIRF100XC; pixel size = 65 nm), was employed. It was housed in a home-built environmental chamber, and all experiments were performed at 37°C. Photobleaching was done for a ≈1 µm diameter circular region for 0.2 s, using 50% of the full laser power. For the photobleaching and recovery observations, another TIRF laser with a 488-nm line (Coherent OBIS 488-100 LS) was used in an oblique illumination mode for the excitation, and the images were recorded with a two-stage microchannel plate intensifier (Hamamatsu Photonics C9016-02MLP24), lens-coupled to an sCMOS camera (Hamamatsu Photonics ORCA-Flash ver4.0 V2 plus C11440-22CU). The images were obtained at a frame rate of 1 Hz, with an exposure time of 33 ms.

### Cell aggregation experiments

L cell clones stably expressing E-cadherin, Nlg1, or Neurexin 1β (Nxn) were grown to 90% confluency in 6 cm cell culture dishes, treated with 1 mL 0.05% trypsin (lacking EDTA; Gibco 15090-046) in Hanks’ balanced salt solution (HBSS; Nissui Pharmaceutical 05906) buffered with 2 mM TES (Dojindo 348-08371), and then incubated at 37°C for 10 min. Cells were detached by mild tapping, and resuspended in the imaging medium. Cells were then centrifuged at 1,000 RPM for 2 min, and the cell pellet was resuspended in the imaging medium. Cells were counted and diluted to approximately 10^6^ cells/mL. The cell suspension (300 µL) was placed in a 1.5 mL Eppendorf tube and used for cell aggregation experiments. When cells from two different clones were mixed (as in the case of Nlg1 + Nxn cell aggregation assays), a 150 µL portion of each cell suspension (at a cell density of 10^6^ cells/mL) was mixed in a 1.5 mL Eppendorf tube. The Eppendorf tubes were immersed in a water-bath at 37°C with shaking at 100 RPM. At each indicated time point, the tube was vortexed briefly to resuspend the cells, and a 10 µL aliquot was placed on the cover glass part of the glass-bottom dish and observed by phase-contrast microscopy. Cell ‘particles’, including both single cells and cell clumps, were counted in the images using a custom ImageJ script. The cell aggregation coefficient was estimated as the number of cell particles detected at the indicated time point divided by the number of cell particles detected at time zero.

### Immunofluorescence

The cultured cells on the FN-coated glass-base dish were fixed with paraformaldehyde (4% w/v; Sigma-Aldrich 158127-500G) at room temperature (RT) for 20 min. The fixed cells were washed twice with HBSS buffered with 2 mM TES, incubated with 0.1% Triton X-100 (Sigma T8787-250ML) and 2% bovine serum albumin (BSA; Sigma A3059-50G) in PBS for 10 min, and then washed twice with imaging medium containing FBS. The cells were immuno-stained for 1 h with an anti-HA primary antibody conjugated to AlexaFluor 555 (Invitrogen 26182-A555) at RT. This was followed by four washes with the imaging medium. The cells were then observed in the imaging medium with the home-built TIRF microscope, which was also used for single molecule observations as described in a previous subsection. Imaging was performed with the 561 nm channel.

### Western blots

Wild-type L cells and L cells transiently expressing PSD95-EGFP were grown to ≈80% confluency in 60-mm plastic dishes, and then disrupted with ice-cold radio immunoprecipitation assay (RIPA) buffer, consisting of 150 mM NaCl, 5 mM ethylenediaminetetraacetic acid (EDTA; Dojindo 345-01865), 1% Nonidet P-40 (Wako 141-08321), 1% sodium deoxycholate (Nacalai Tesque 08805-72), 0.1% sodium dodecyl sulfate (SDS, Wako 191-07145), and 100x diluted protease inhibitor cocktail (Nacalai Tesque 25955-24), buffered with 50 mM Tris-HCl at pH 7.4 (350 µL), and kept on ice for 5 min. The entire material in the dish was collected by using a cell scraper, and then centrifuged at 15,000 rpm for 20 min. The supernatant (300 µL) was mixed with 100 µL of 4x Laemmli sample buffer, consisting of 8% SDS, 0.12% bromophenol blue, 40% glycerol, and 400 mM dithiothreitol, buffered with 200 mM Tris-HCl at pH 6.8, and boiled for 5 min. The boiled mixture (2-10 µL, 1-2 µg of total protein for each lane) was loaded on a precast 4-20% gradient polyacrylamide gel (PAGE, Bio-Rad 4561096) and electrophoresed at a constant voltage of 200 V for 35 min using a Mini Protean Tetra Cell (Bio-Rad 1658000JA), in Tris-glycine running buffer (25 mM Tris-HCl, 190 mM glycine, and 0.1% SDS).

The separated proteins were blotted onto polyvinylidene fluoride (PVDF) membranes by the semi-dry transfer method, using a Trans-Blot Turbo (Bio-Rad 1704150, mixed MW protocol) with the PVDF transfer pack (Bio-Rad 1704156). The immuno-detection was performed by a SNAP i.d. (Millipore SNAP2BASE), according to the manufacturer’s recommendations. The membranes were blocked with Blocking One (Nacalai Tesque 03953-95) and washed with Tris-buffered saline (TBS; 137 mM NaCl and 2.7 mM KCl buffered with 25 mM Tris-HCl, pH 7.4) supplemented with 0.1% Tween 20 (TBS-T; Sigma P7949-500ML).

Primary antibodies were as follows: mouse anti-PSD95 (Synaptic Systems 124 011, Lot#124 011/1-21, 1 µg/mL) and mouse anti-GAPDH (ProteinTech 60004-1-Ig, Lot#10008047, 1 µg/mL). The secondary antibody IgG used was horseradish peroxidase (HRP)-conjugated goat anti-mouse immunoglobulin G (IgG, Invitrogen G21040, Lot#2122350, 0.5 mg/mL).

The membrane was incubated with Western BLoT Hyper HRP Substrate (Takara T7103B) for chemiluminescence detection, and then imaged with a Fuji Film LAS-3000 using the high sensitivity mode or a Thermo Fischer Scientific iBright FL1500 image analyzer (2x2 binning, 1.5x zoom).

### Statistical analysis and software

Conventional TIRF images were processed and analyzed using ImageJ (Schindelin et al., 2012), and ilastik (Berg et al., 2019). Curve fittings were performed using Origin 2016 (OriginLab).

Statistical analyses were performed using OriginPro 2016 and Excel (Microsoft). P<0.05 was defined as statistically significant. Figures and videos were edited using BioRender (biorender.com) and ImageJ, respectively. No statistical methods were used to determine sample sizes prior to the experiments. Sample sizes were comparable to many studies using similar experimental approaches.

### Figures

Figures were created using BioRender, and can be accessed at the following link. Acharya, S. (2025) https://BioRender.com/p0bi90l

### Data availability

All data (including raw data and all analyses) can be made available upon reasonable request.

## SUPPLEMENTARY FIGURE LEGENDS

**Supplementary Figure 1.**
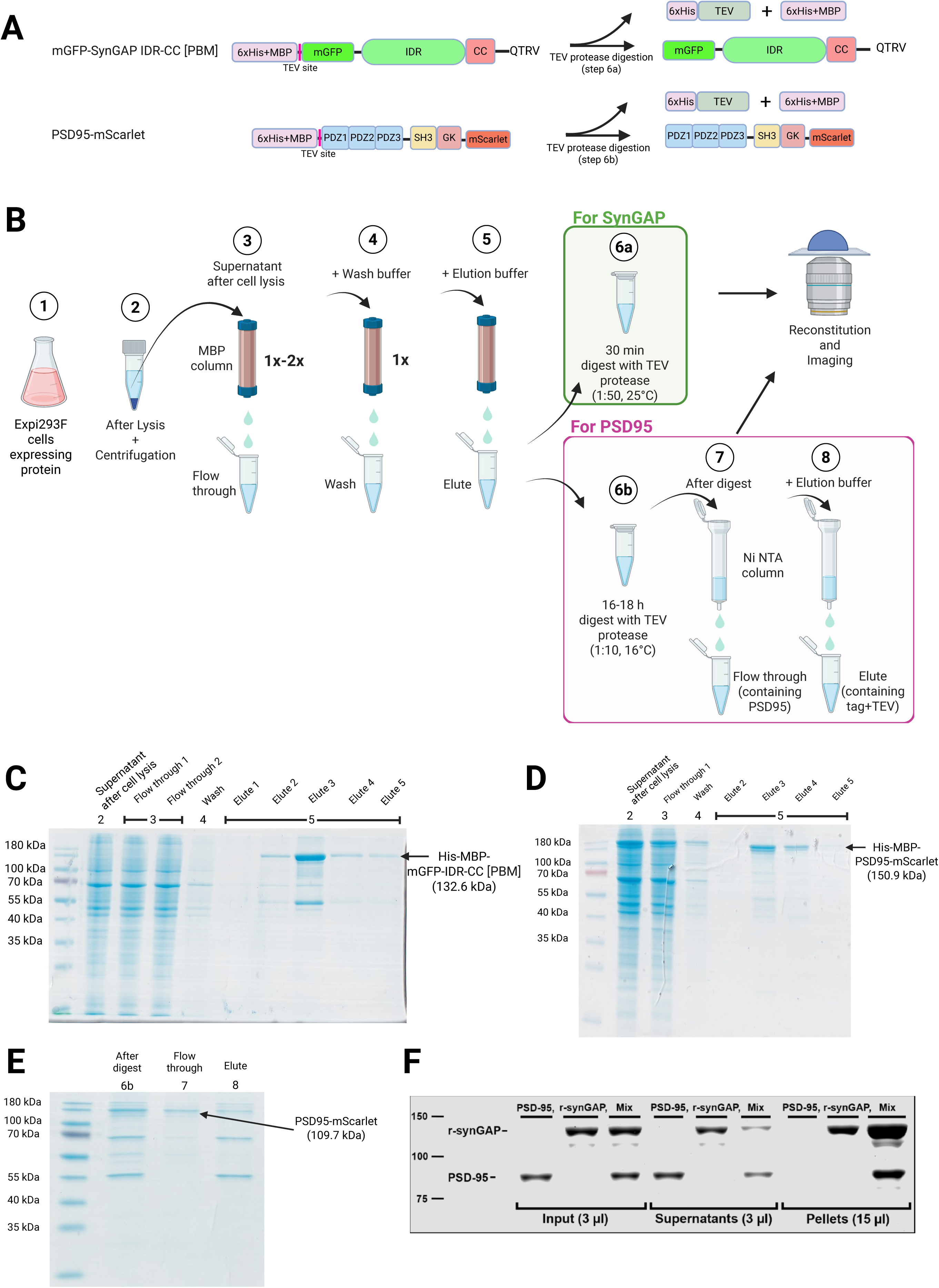
(A) Schematics of mGFP-(SynGAPα1)-IDR-CC[PBM] and PSD95-mScarlet proteins isolated by affinity chromatography. Both constructs contained the N-terminal MBP tag (including 6x His sequence) followed by a TEV protease site. For both proteins, the MBP tag was removed by an incubation with TEV protease, either during the *in-vitro* reconstitution for mGFP-(SynGAPα1)-IDR-CC[PBM] before observation (**step 6a in Supplementary Figure 1B**) or as a part of the purification protocol for PSD95-mScarlet (**step 6b in Supplementary Figure 1B**). (B) Schematic figure showing the key steps for purifying the proteins shown in **A**. At Step 6b for PSD95-mScarlet, the His-MBP affinity tag was cleaved by His-TEV protease, and the cleaved MBP tag and TEV protease were removed by the Nickel affinity chromatography (flowthrough at Step 7 contains PSD95-mScarlet). At Step 6a for mGFP-(SynGAPα1)-IDR-CC[PBM], the His-MBP affinity tag was cleaved just prior to the *in-vitro* reconstitution assays by an incubation with TEV protease for 30 min. **(C-E)** SDS-PAGE gels stained with Coomassie Brilliant Blue, showing the purification steps of mGFP-(SynGAPα1)-IDR-CC[PBM] **(C)** and PSD95-mScarlet **(D, E)**. The numbers on the lanes indicate the purification steps shown in **B**. (F) Three test-tube condensation assays: 1.5 µM unlabeled PSD95 (PSD95), 1.5 µM unlabeled r-SynGAP (r-synGAP), and 1.5 µM each of PSD95 and r-synGAP (Mix) in standard buffer at RT. Immediately after the addition of protein to the standard buffer, aliquots of inputs were removed and mixed with SDS-gel sample buffer (Input). After 1 min, the three assay tubes were centrifuged at 16,000g for 10 min at RT. Supernatants were removed and added to SDS-gel sample buffer (Supernatants). Pellets were resuspended in SDS-gel sample buffer (Pellets). Samples were fractionated by SDS-PAGE. Proteins were visualized with Gel-Code Blue, and scanned in a LICOR Odyssey imager, which digitally records density over six logs of dynamic range. Protein in each band was quantified with the LICOR software by comparison to BSA standards, and corrected to report equivalent volumes. The PSD95 supernatant contained 99±0.2% and the pellet 1±0.2%; synGAP 77±1.3% and 23±1.3%, respectively. The Mix supernatant contained 67±1.5% and the pellet 33±1.5% of the total PSD95. The Mix supernatant contained 41±1.6% and the pellet 59±1.6% of the total synGAP. Errors are SE.

**Supplementary Figure 2.**
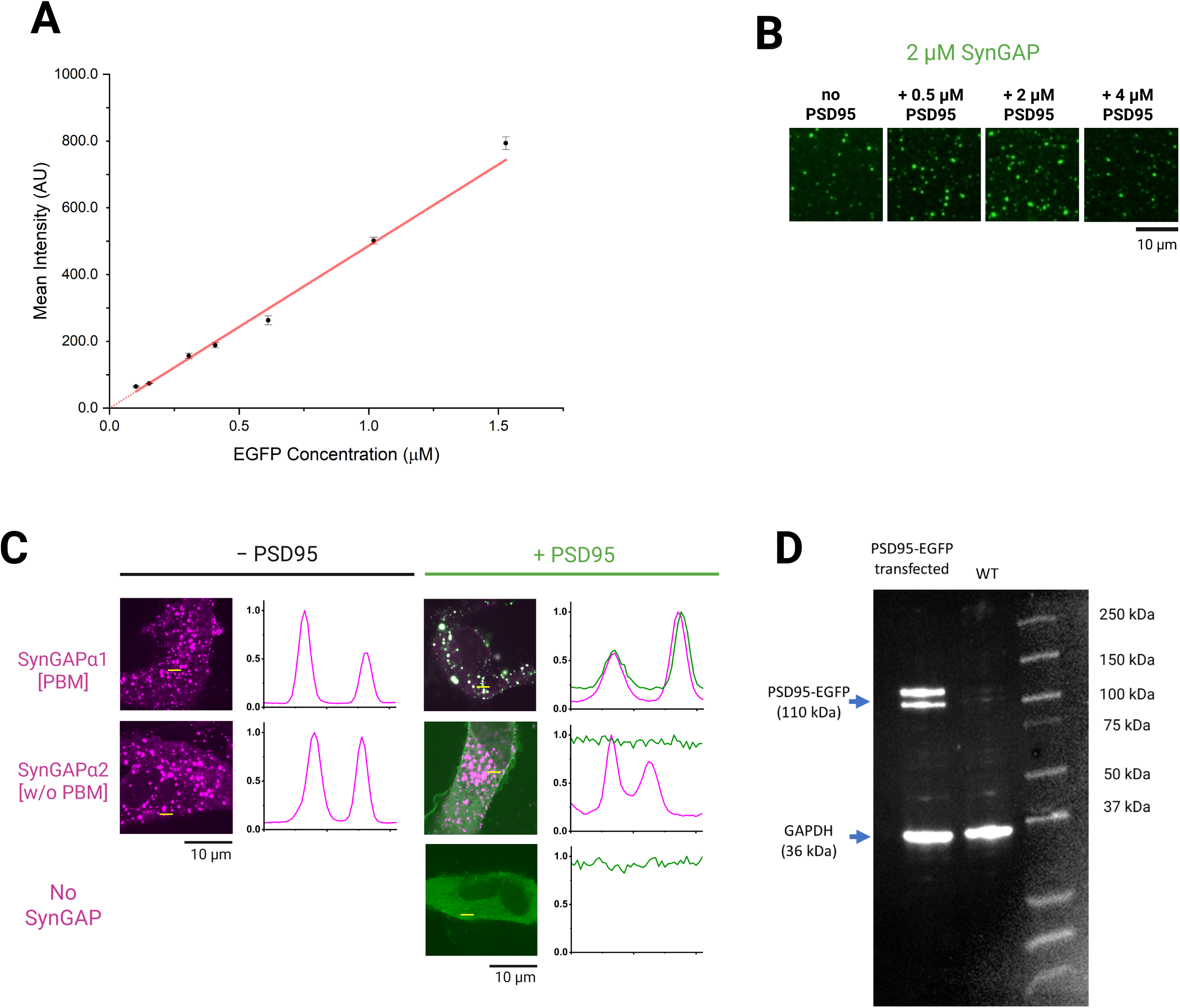
(A) Calibration of the EGFP concentration based on the fluorescence intensity observed by confocal microscopy. Purified EGFP (Biovision 499-100) in the imaging buffer was imaged at 2 µm above the glass using confocal microscopy. Dots, mean intensity values; bars, SEMs (3 independent measurements). The data points were fitted with a linear function. (B) Representative images of 2 µM mGFP-(SynGAPα1)-IDR-CC[PBM] as a function of the PSD95-mScarlet concentration (not visualized) 2 min after 1% PEG8000 addition, suggesting that mGFP-(SynGAPα1)-IDR-CC[PBM] condensation might be enhanced in the presence of approximately equimolar concentrations of PSD95. (C) Another set of representative images and line scans as shown in Fig. 2C, but with the use of live HeLa cells rather than L cells. Here, the intensity profile of the line scan is normalized by the maximal intensity of each line scan. (D) L cells do not express detectable amounts of PSD95. The western blot was probed with anti-PSD95 polyclonal antibodies.

**Supplementary Figure 3.**
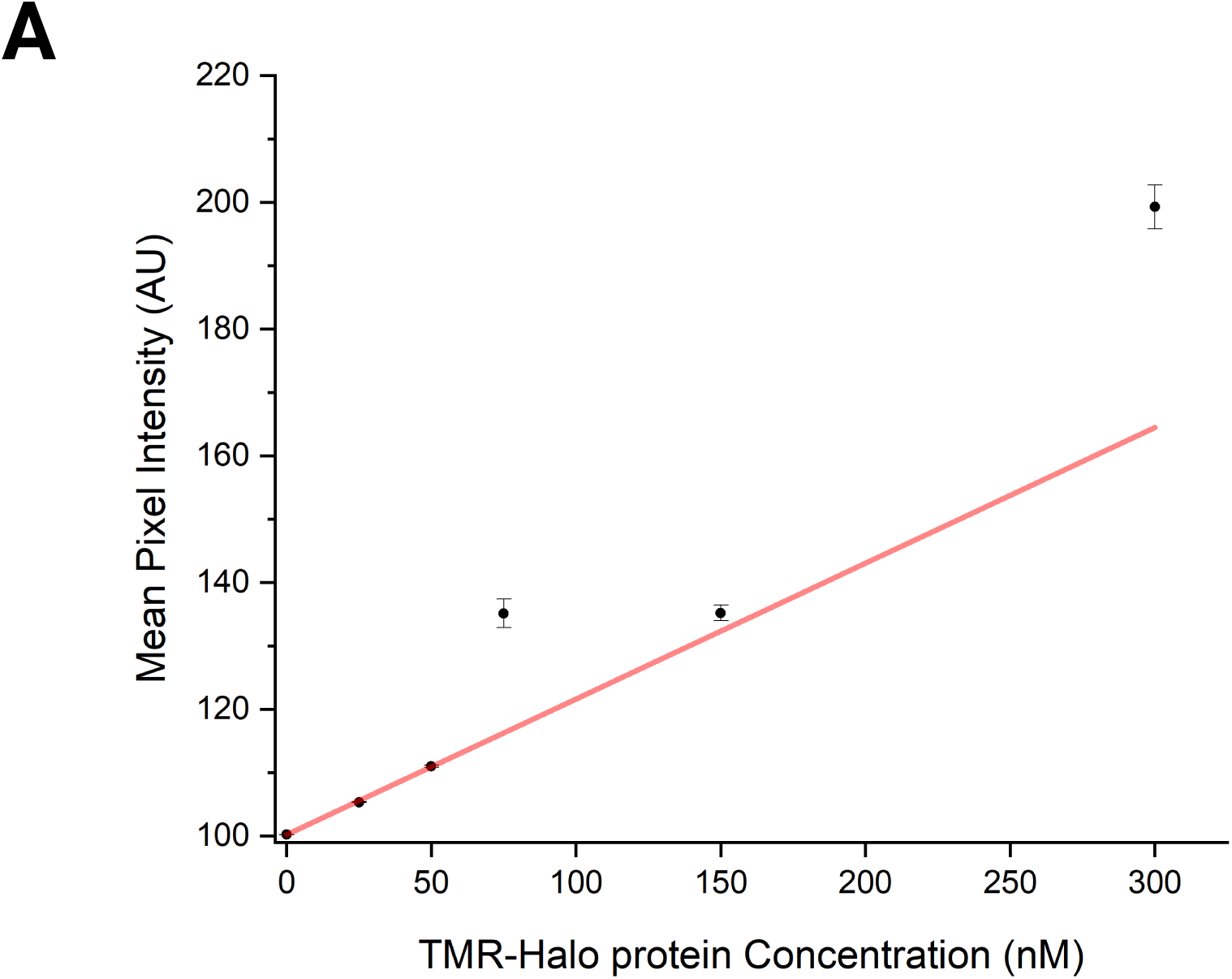
(A) Calibration of the TMR labelled Halo protein concentration based on the fluorescence intensity observed by confocal microscopy. Purified Halo protein (Promega G4491; labelled with TMR-Halo ligand), in the imaging buffer was imaged at 2 µm above the glass using confocal microscopy. Dots, mean intensity values; bars, SEMs (3 independent measurements). The data points were fitted with a linear function.

**Supplementary Figure 4.**
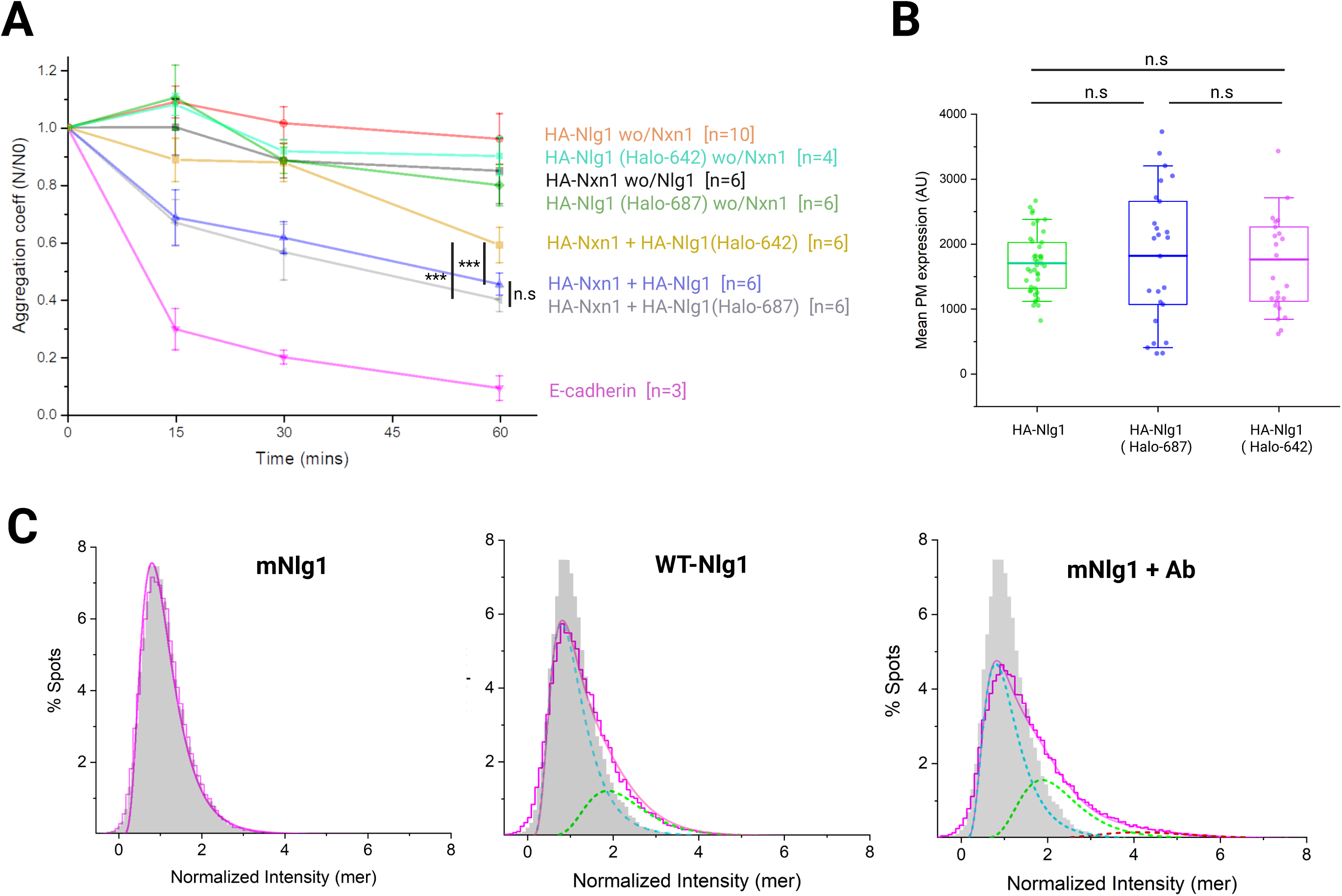
(A) Adhesion of HA-tagged Nlg1 (HA-Nlg1) and HA-Nlg1 linked to a Halo7 tag at amino acid 642 or 687 were tested in a cell-cell aggregation assay for examining the adhesion functions of Halo-tagged Nlg1. For the visualization of Nlg1 expression in cells, Nlg1 and Halo-Nlg1 contained HA tags at their N-termini. Previously, the GFP11 fragment was inserted at amino acid 642 (Tsetsenis et al., 2014), but a new tag position at amino acid 687 was also tested. L cells stably expressing the test HA-Nlg1 constructs were mixed with L cells stably expressing HA-tagged neurexin1β (HA-Nxn) or L cells without HA-Nxn expression (control). As a positive control, L cells expressing E-cadherin, which undergoes homophilic adhesion, were employed. The progress of cell aggregation was monitored by evaluating the aggregation coefficient, which was defined as the number of particles (single cells + cell clusters) at time t divided by that at time 0 (see Methods for details). Dots and bars indicate the means and SEMs. ‘n’ values indicate the number of replicates. Asterisks and n.s. indicate *p* < and > 0.05, respectively, using the one-way ANOVA test. (B) In the cell aggregation assay shown in A, we selected the L cells stably expressing about the same number of HA-Nlg1, HA-Nlg1 (Halo-687), and HA-Nlg1 (Halo-642) molecules in the PM, as shown here. The expression levels were assessed by monitoring the fluorescence intensity of the live L cells 1 h after the addition of Alexa-555-conjugated anti-HA antibodies. HA-Nlg1, 41 cells from 5 dishes; HA-Nlg1 (Halo-687), 25 cells from 5 dishes; and HA-Nlg1 (Halo-642), 25 cells from 5 dishes. Bars, boxes, and whiskers indicate the mean values, inter-quartile ranges (25% - 75%), and 10% - 90% ranges, respectively. n.s. indicates *p* > 0.05, using the one-way ANOVA test. (C) Basic data to determine the number fractions of molecules existing as monomers, dimers, trimers, and tetramers (Fig. 5A). The distributions of the signal intensities of single individual fluorescent spots of TMR bound to Halo7 (for Nlg1s, magenta histograms) and the Halo7-tagged transmembrane domain of the LDL receptor, which is a monomer reference non-raft molecule (Halo7-TM; grey histograms, which are the same for the three histograms, n = 32,891 spots in 17 cells). First, the histogram of TMR-Halo7-TM (the same for all 3 graphs; n = 32,891 spots in 17 cells) was fitted with a single lognormal function, with a mode of 58,402 and log standard deviation of 0.48. These values were used for producing the lognormal functions of dimers, trimers, and tetramers, and the histograms of other molecules were fitted with the sum of these lognormal functions using their pre-factors as fitting parameters (Morise et al., 2019; smooth magenta curves). The distribution for mNlg1 (n = 41,785 spots from 19 cells) was virtually the same as that for Halo7-TM, and could be best fitted with a single lognormal function, indicating that these molecules exist as monomers. The distributions for WT-Nlg1 (n = 30,573 spots from 15 cells) and mNlg1+Ab (n = 55,652 spots from 20 cells) were fitted with the sum of four lognormal functions (dashed curves in the graphs; monomers in cyan, dimers in green, trimers in orange, and tetramers in red).

**Supplementary Figure 5.**
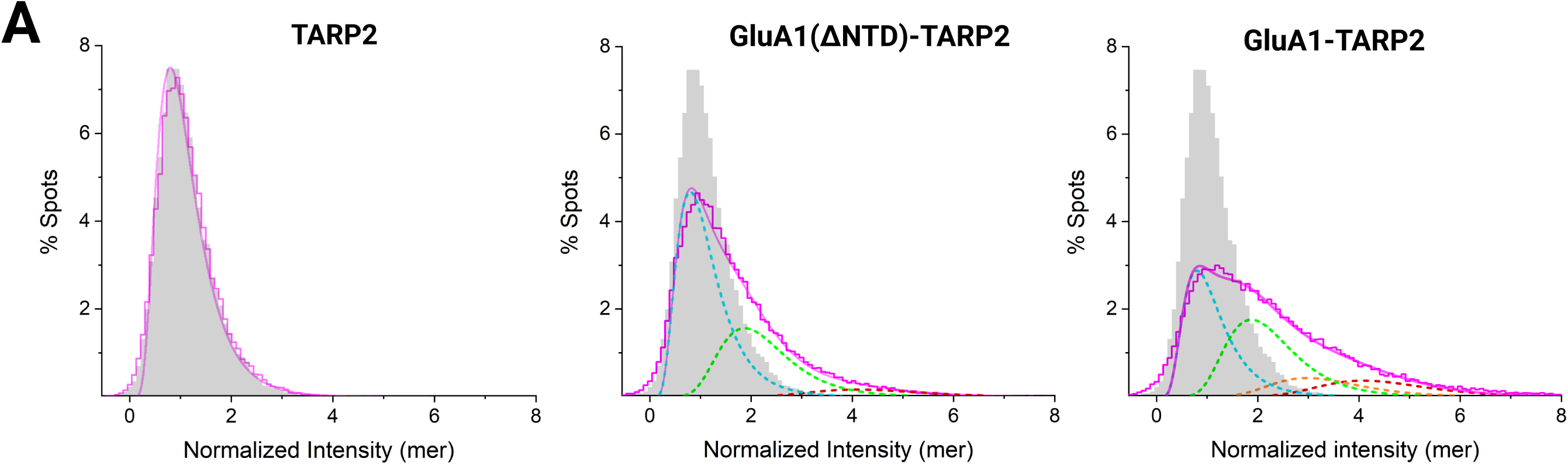
(A) Basic data to determine the number fractions of molecules existing as monomers, dimers, trimers, and tetramers (Fig. 6A). Halo7-TM data are the same as those shown in **Supplementary Fig. 4C**. The distribution for TARP2 (n = 29,132 spots from 17 cells) is virtually the same as that for Halo7-TM. The distributions for TARP2-GluA1(ΔNTD) (n = 42,065 spots from 25 cells) and TARP2-GluA1 (n = 74,888 spots from 22 cells) are displayed in the same manner as in **Supplementary Fig. 4C**.

## Notes

### Competing Interest Statement

The authors have declared no competing interest.

