## Supplementary Tables for "SynGAP forms biocondensates at sub-micromolar concentrations and recruits PSD95 and receptor oligomers, functioning as a key initiator of PSD formation"

**Table 1. Summary of number of immobilization events and cells analyzed for various Nlg molecules**

Immobilization events of mNlg1, WT-Nlg1, mNlg1+Ab single-molecule spots (from 8, 7, and 12 independent movies) were analyzed.

| **Molecule** | **Immobilization events** | **Cells sampled** |
| --- | --- | --- |
| mNlg1 | 11,571 | 8 |
| WT-Nlg1 | 12,671 | 7 |
| mNlg1+Ab | 8,614 | 12 |

**Table 2. Summary of number of immobilization events and cells analyzed for various AMPA molecules**

Immobilization events of TARP2, GluA1(ΔNTD)-TARP2, GluA1-TARP2 spots (from 13, 9, and 13 independent movies) were analyzed.

| **Molecule** | **Immobilization events** | **Cells sampled** |
| --- | --- | --- |
| TARP2 | 16,142 | 13 |
| GluA1(ΔNTD)-TARP2 | 9,981 | 9 |
| GluA1-TARP2 | 10,752 | 13 |

**Table 3. Statistical analysis of the distributions of immobilization durations of various Nlg molecules.**

| **Test** | ***P* value** |
| --- | --- |
| Kruskal-Wallis multiple comparison test (WT-Nlg1, mNlg1, mNlg1+Ab) | < 2.2 × 10^-16^ |
| Steel-Dwass pair tests |  |
| mNlg1 vs WT-Nlg1 | < 2.2 × 10^-8^ |
| mNlg1 vs mNlg1+Ab | < 2.2 × 10^-8^ |
| WT-Nlg1 vs mNlg1+Ab | < 2.2 × 10^-8^ |

**Table 4. Statistical analysis of the distributions of immobilization durations of various AMPA molecules.**

| **Test** | ***P* value** |
| --- | --- |
| Kruskal-Wallis multiple comparison test (TARP2, GluA1(ΔNTD)-TARP2, GluA1-TARP2) | < 2.2 × 10^-16^ |
| Steel-Dwass pair tests |  |
| TARP2 vs GluA1(ΔNTD)-TARP2 | 0.0022 |
| TARP2 vs GluA1-TARP2 | < 2.2 × 10^-8^ |
| GluA1(ΔNTD)-TARP2 vs GluA1-TARP2 | < 2.2 × 10^-8^ |

**Table 5. Statistical analysis of clustering coefficient vs concentration curves of various SynGAP molecules shown in Fig. 2A, in the concentration range of 0 – 0.25 µM**

| **Test** | ***P* value** |
| --- | --- |
| ANCOVA multiple comparison test |  |
| α1 vs α2 | 0.9999 |
| α1 vs IDR-CC-PBM | 0.8737 |
| α1 vs ΔIDR | 7.3679 × 10^-8^ |
| α1 vs ΔCC-PBM | 1.1484 × 10^-9^ |
| α1 vs IDR only | 1.4212 × 10^-11^ |
| α1 vs CC-PBM only | 2.9323 × 10^-11^ |
| α2 vs IDR-CC-PBM | 0.9490 |
| α2 vs ΔIDR | 0.0004 |
| α2 vs ΔCC-PBM | 2.9531 × 10^-11^ |
| α2 vs IDR only | 5.5207 × 10^-7^ |
| α2 vs CC-PBM only | 7.3513 × 10^-7^ |
| IDR-CC-PBM vs ΔIDR | 0.0028 |
| IDR-CC-PBM vs ΔCC-PBM | 1.6551 × 10^-5^ |
| IDR-CC-PBM vs IDR only | 2.0877 × 10^-6^ |
| IDR-CC-PBM vs CC-PBM only | 2.9793 × 10^-6^ |
| ΔIDR vs ΔCC-PBM | 0.2158 |
| ΔIDR vs IDR only | 0.1062 |
| ΔIDR vs CC-PBM only | 0.1234 |
| ΔCC-PBM vs IDR only | 0.9999 |
| ΔCC-PBM vs CC-PBM only | 0.9999 |
| IDR only vs CC-PBM only | 0.9999 |

**Table 6. Statistical analysis of clustering coefficient vs concentration curves of various SynGAP molecules shown in Fig. 2A, in the concentration range of > 0.3 µM**

| **Test** | ***P* value** |
| --- | --- |
| ANCOVA multiple comparison test |  |
| α1 vs α2 | 0.9672 |
| α1 vs IDR-CC-PBM | 0.9971 |
| α1 vs ΔIDR | 0.9899 |
| α1 vs ΔCC-PBM | 1.1038 × 10^-5^ |
| α1 vs IDR only | 1.6846 × 10^-15^ |
| α1 vs CC-PBM only | 7.2250 × 10^-12^ |
| α2 vs IDR-CC-PBM | 0.9998 |
| α2 vs ΔIDR | 0.9998 |
| α2 vs ΔCC-PBM | 0.0122 |
| α2 vs IDR only | 1.1609 × 10^-8^ |
| α2 vs CC-PBM only | 4.0443 × 10^-7^ |
| IDR-CC-PBM vs ΔIDR | 0.9999 |
| IDR-CC-PBM vs ΔCC-PBM | 0.0007 |
| IDR-CC-PBM vs IDR only | 1.4117 × 10^-11^ |
| IDR-CC-PBM vs CC-PBM only | 3.4874 × 10^-9^ |
| ΔIDR vs ΔCC-PBM | 4.4087 × 10^-5^ |
| ΔIDR vs IDR only | 1.4873 × 10^-15^ |
| ΔIDR vs CC-PBM only | 2.0809 × 10^-11^ |
| ΔCC-PBM vs IDR only | 0.0179 |
| ΔCC-PBM vs CC-PBM only | 0.0594 |
| IDR only vs CC-PBM only | 0.9999 |

**Table 7. Statistical analysis of clustering coefficient vs concentration curves of various SynGAP molecules shown in Fig. 2B, in the concentration range of 0 – 0.25 µM**

| **Test** | ***P* value** |
| --- | --- |
| ANCOVA multiple comparison test |  |
| α1 vs α1 + CaMKII Kinase dead (K42M) | 0.1089 |
| α1 vs α1 + CaMKII Constitutively active (T286D) | 7.1419 × 10^-7^ |
| α1 vs α1 Phosphomimetic mutant | 3.8921 × 10^-5^ |
| α1 + CaMKII Kinase dead (K42M) vs α1 + CaMKII Constitutively active (T286D) | 0.0040 |
| α1 + CaMKII Kinase dead (K42M) vs α1 Phosphomimetic mutant | 0.0502 |
| α1 + CaMKII Constitutively active (T286D) vs α1 Phosphomimetic mutant | 0.8915 |

**Table 8. Statistical analysis of clustering coefficient vs concentration curves of various SynGAP molecules shown in Fig. 2B, in the concentration range of >0.3 µM**

| **Test** | ***P* value** |
| --- | --- |
| ANCOVA multiple comparison test |  |
| α1 vs α1 + CaMKII Kinase dead (K42M) | 0.9999 |
| α1 vs α1 + CaMKII Constitutively active (T286D) | 0.7903 |
| α1 vs α1 Phosphomimetic mutant | 0.3335 |
| α1 + CaMKII Kinase dead (K42M) vs α1 + CaMKII Constitutively active (T286D) | 0.8218 |
| α1 + CaMKII Kinase dead (K42M) vs α1 Phosphomimetic mutant | 0.4227 |
| α1 + CaMKII Constitutively active (T286D) vs α1 Phosphomimetic mutant | 0.8881 |
